# Chromosomal fusions and large-scale inversions are key features for adaptation in Arctic codfish species

**DOI:** 10.1101/2024.06.28.599280

**Authors:** Siv N.K Hoff, Marius Maurstad, Ole K. Tørresen, Paul R. Berg, Kim Præbel, Kjetill S. Jakobsen, Sissel Jentoft

## Abstract

The evolutionary impact of structural variants, such as chromosomal inversions, is well documented, especially for their role in local adaptation in high gene flow systems. However, the role of other genomic rearrangements like chromosomal fusions, fissions, and translocations is still relatively unexplored. Here we present six chromosome-level Gadid reference genomes for the non-migratory Atlantic cod (*Gadus morhua*) i.e., Norwegian coastal cod (NCC), Atlantic haddock *(Melanogrammus aeglefinus),* burbot (*Lota lota*), European hake (*Merluccius merluccius*) as well as two keystone Arctic codfishes: the polar cod (*Boreogadus saida*) and Arctic cod (*Arctogadus glacialis*). Within a comparative genomics framework, we uncovered several lineage-specific chromosomal fusions, resulting in a reduced number of chromosomes compared to the ancestral state in the two cold-water adapted codfishes. The identified fusions were not homologous, i.e., indicating that they originate from independent evolutionary events. Additionally, a high number of partly overlapping chromosomal inversions between the two species were detected. Using a smaller population dataset, we uncovered a high degree of conservation for some of the overlapping inversions (including some breakpoint regions), suggesting that these regions are under selection, and potentially of evolutionary importance. With the use of chromosome-level genome assemblies, we demonstrate how large genomic reorganizations are likely to play important roles in speciation processes and thus, in particular to adaptation to freezing environmental conditions. Moreover, we observe that such massive rearrangement events can take place across relatively short evolutionary time scales.

## Background

Genomic architecture and organization — including chromosome numbers — vary massively across the tree of life[1,2]. Most striking variation is reported within the Plantae kingdom[1,2], mainly due to whole-genome duplications and/or polyploidization events as well as via single chromosome changes such as chromosomal fusions and fissions[1,2]. For closely related species, genome organization has in general been thought to be relatively conserved[3,4]. However, in parallel with the developments in high throughput sequencing technologies, several studies have demonstrated how chromosomal reorganizations can occur over relatively short evolutionary timescales[5] and in some cases lead to chromosomal speciation (reviewed in Damas et al.[6]) where examples include blind mole rats[7], rock-wallaby[8] and cichlid fish[9]. Large-scale genomic rearrangements, including chromosomal inversions, translocations, fission, and fusions can be of high evolutionary importance — by linking allelic variants and/or genetic elements — leading to alterations in morphological or behavioral traits[10–13]. Ultimately, such genomic reorganizations can manifest within species, populations and/or ecotypes as genetic polymorphisms, in which natural selection can act upon, and mediate genomic divergence between species, populations, or ecotypes.

Teleostei is the most diverse and species-rich infraclass of vertebrates with more than 30,000 species inhabiting numerous marine and freshwater habitats[14]. Not surprisingly, pronounced genomic plasticity has been reported, encompassing variations in genome organization, size, and chromosome numbers, all features that could be instrumental for the remarkable diversity observed within this taxon[15–18]. In general, a higher number of chromosomes as well as increased karyotype variation have been observed in freshwater species compared to their marine counterparts[19,20]. This feature is thought to be linked to the higher number of unique habitats, with well-defined barriers and lack of gene flow, as well as greater variability in environmental conditions encountered in freshwater vs. marine habitats[19,20]. For instance, species within the genus *Nothobranchius* — adapted to the fluctuating environments of East African temporal savannah pools — display extensive karyotype variability and the diploid chromosome numbers range from 16 to 50[20]. While genomic organization and chromosomal numbers tend to be more stable in marine species, some exceptions are reported. For the suborder Notothenioidei, an endemic group adapted to the freezing conditions of the Southern Oceans — multiple lineages have undergone extensive chromosomal reorganizations among closely related species[21–24]. Specifically, within the genera *Trematomus*[22] and *Notothenia*[23] unique chromosomal fusions and fissions have been reported, which resulted in reduced chromosome numbers for several of the species.

In the present study, we take a close look at the northern gadids (codfishes), belonging to a species-rich family within the order *Gadiformes,* which includes species primarily distributed across the northern Atlantic, including the freezing Arctic Oceans and the Northern Pacific[25]. Although Arctic conditions are not as extreme as in Antarctica, organisms in both regions face similar environmental challenges, such as freezing seawater and thick ice covers[26]. For instance, similar to the notothenioids, cold water adapted gadids such as the polar cod (*Boreogadus saida*) and Arctic cod (*Arctogadus glacialis*) display an extended repertoire of antifreeze glycoproteins compared to their temperate counterparts, i.e., enabling these species to cope under extreme freezing conditions[27,28]. Moreover, inter- and intraspecies chromosome number variation has been reported in previous studies within the gadid family (see Figure 1 and Table S1). Among the cold-water adapted species residing in the Arctic and sub-Arctic regions[29,30] fewer chromosomes have been identified compared to the presumed ancestral teleost karyotype (n=24-26)[31–33]. The reduction in chromosome number seen both in the Antarctic notothenioids and Arctic codfishes is intriguing and raises questions about the adaptive significance of chromosomal fusions in species inhabiting high-latitude oceans. For Arctic cod observed population-level karyotype variation seems to follow a latitudinal cline, supporting the idea of chromosomal reduction being advantageous in cold-water conditions[30]. Moreover, high numbers of chromosomal inversions have been reported for many gadids, and are likely to play important roles in the separation of co-occurring cryptic ecotypes[34,35], as well as for local adaptation to different environmental conditions[36–39].

**Figure 1:**
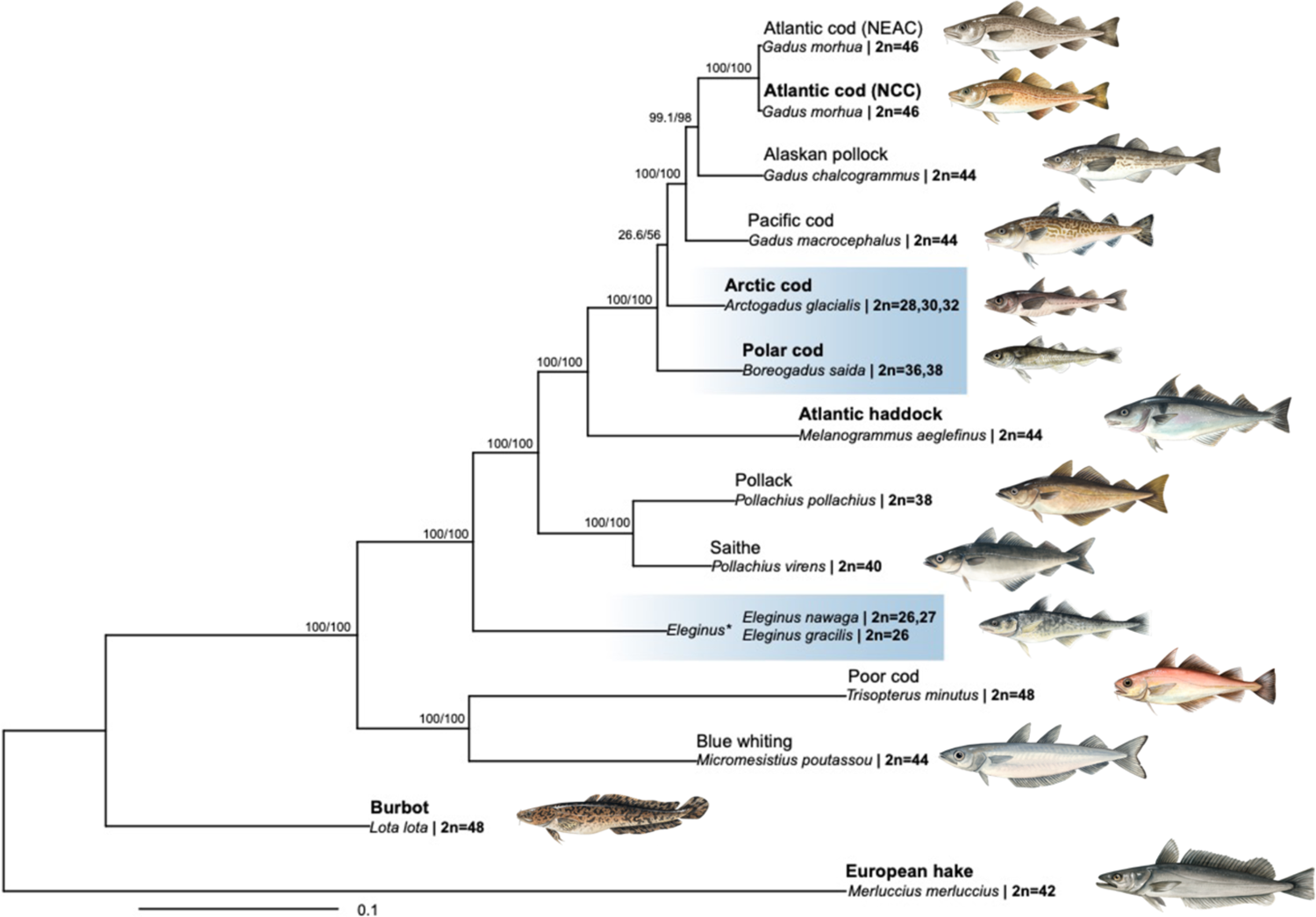
Karyotype variability and phylogenetic relationships among codfishes. Overview of the chromosome numbers in 13 Gadiform fish species (see Table S1 for more details) shown with their phylogenetic relationship generated by IQ-TREE2 v2.2.0[132], using the 13 mitochondrial protein-coding genes. Support is given as SH-alrt/UFBoot (see Materials and Methods for details). The six species genome sequenced in the present study are marked in bold. The Arctic residing codfishes are marked in blue. *=*Eleginus gracilis* here used as a representative for Saffron cod (*Eleginus gracilis*, 2n=26) and Navaga (*Eleginus nawaga*, 2n=26-27). Illustrations made by Alexandra Viertler.

To further explore the evolutionary genomic alterations and reorganizations that have taken place within the gadids, and how these modifications could have impacted the inference of their phylogenetic relationship, we have generated and taken advantage of chromosome-level genome assemblies for six Gadiform species: Arctic cod (*Arctogadus glacialis*), polar cod (*Boreogadus saida*), non-migratory Atlantic cod (*Gadus morhua*) i.e., Norwegian coastal cod (NCC), Atlantic haddock *(Melanogrammus aeglefinus),* burbot (*Lota lota*) and European hake (*Merluccius merluccius*). By utilizing a comparative genomic approach — combined with population data from Arctic cod and polar cod — we identify several lineage-specific chromosomal fusions as well as a high number of partly overlapping chromosomal inversions present in the two cold-water adapted codfishes. Intriguingly, high degree of conservation was detected for some of the overlapping inversions as well as for the breakpoint regions, suggesting that these regions are under strong directional and/or purifying selection, and potentially of high evolutionary impact linking genes of importance for local adaptation. Lastly, we reconstruct phylogenetic trees using Benchmarking Universal Single-Copy Orthologs (BUSCO) and mitochondrial genes in an effort to elucidate the relationship of Arctic cod within the Gadidae family.

## Results

### Generation of chromosome-anchored genome assemblies for six codfish species

The generation of primary assemblies using long-read PacBio data ranged from 42.42 to 98.97 Gb of data (see Table S2 for more information) and subsequently scaffolding with the linked read data, e.g., 10X and/or Illumina Hi-C, yielded highly contiguous chromosome-level genome assemblies for all six species: polar cod, Arctic cod, Norwegian coastal cod, Atlantic haddock, burbot, and European hake. The number of super scaffolds obtained from each assembly utilizing Hi-C data matched the karyotype numbers earlier reported for five of the species (Table 1, Table S1, and Figure S1A-F for karyotype overview), except the genome assembly of burbot which resulted in 23 chromosome length scaffolds (n=24 reported cytogenetically[40,41]). The genome assemblies ranged from ∼538 Mb across 2223 to ∼679 Mb across 4087 scaffolds for burbot and Atlantic cod (NCC), respectively (Table 1). All putative chromosomes assembled for the six species were larger than 10 Mb in size, and the majority of the assembled sequences (> 93.13% for all assemblies) were placed within the chromosome-length scaffolds (Table 1). Gene completeness was scored to > 90.1% (complete BUSCOs) for the genome assemblies (Table 1).

**Table 1.**
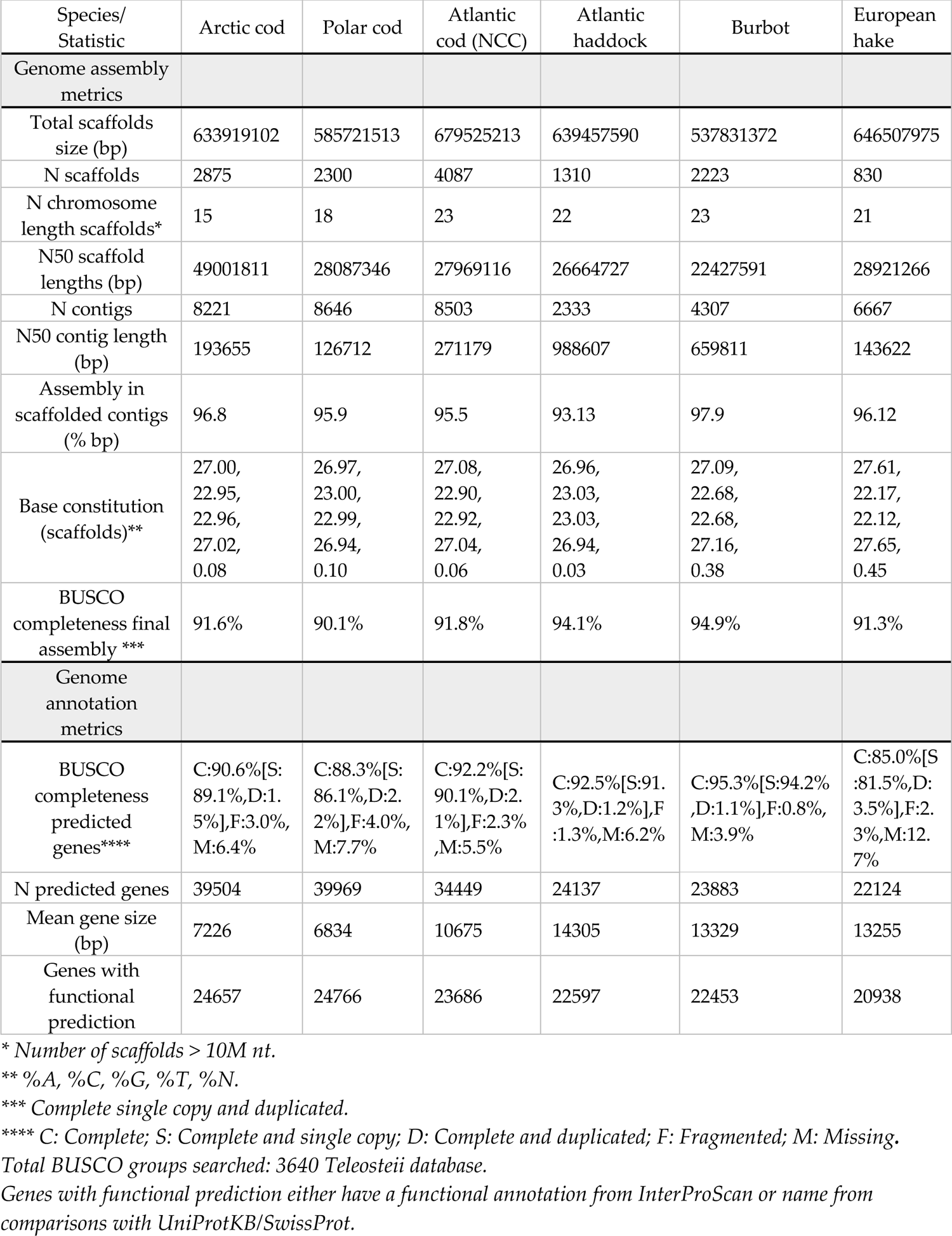
Assembly metrics of final genome assemblies and annotation for the six species.

| Species/<br>Statistic | Arctic cod | Polar cod | Atlantic<br>cod (NCC) | Atlantic<br>haddock | Burbot | European<br>hake |
| --- | --- | --- | --- | --- | --- | --- |
| Genome assembly<br>metrics |  |  |  |  |  |  |
| Total scaffolds<br>size (bp) | 633919102 | 585721513 | 679525213 | 639457590 | 537831372 | 646507975 |
| N scaffolds | 2875 | 2300 | 4087 | 1310 | 2223 | 830 |
| N chromosome<br>length scaffolds* | 15 | 18 | 23 | 22 | 23 | 21 |
| N50 scaffold<br>lengths (bp) | 49001811 | 28087346 | 27969116 | 26664727 | 22427591 | 28921266 |
| N contigs | 8221 | 8646 | 8503 | 2333 | 4307 | 6667 |
| N50 contig length<br>(bp) | 193655 | 126712 | 271179 | 988607 | 659811 | 143622 |
| Assembly in<br>scaffolded contigs<br>(% bp) | 96.8 | 95.9 | 95.5 | 93.13 | 97.9 | 96.12 |
| Base constitution<br>(scaffolds)** | 27.00,<br>22.95,<br>22.96,<br>27.02,<br>0.08 | 26.97,<br>23.00,<br>22.99,<br>26.94,<br>0.10 | 27.08,<br>22.90,<br>22.92,<br>27.04,<br>0.06 | 26.96,<br>23.03,<br>23.03,<br>26.94,<br>0.03 | 27.09,<br>22.68,<br>22.68,<br>27.16,<br>0.38 | 27.61,<br>22.17,<br>22.12,<br>27.65,<br>0.45 |
| BUSCO<br>completeness final<br>assembly *** | 91.6% | 90.1% | 91.8% | 94.1% | 94.9% | 91.3% |
| Genome<br>annotation<br>metrics |  |  |  |  |  |  |
| BUSCO<br>completeness<br>predicted<br>genes**** | C:90.6%[S:<br>89.1%,D:1.<br>5%],F:3.0%,<br>M:6.4% | C:88.3%[S:<br>86.1%,D:2.<br>2%],F:4.0%,<br>M:7.7% | C:92.2%[S:<br>90.1%,D:2.<br>1%],F:2.3%<br>,M:5.5% | C:92.5%[S:91.<br>3%,D:1.2%],F<br>:1.3%,M:6.2% | C:95.3%[S:94.2%<br>,D:1.1%],F:0.8%,<br>M:3.9% | C:85.0%[S<br>:81.5%,D:<br>3.5%],F:2.<br>3%,M:12.<br>7% |
| N predicted genes | 39504 | 39969 | 34449 | 24137 | 23883 | 22124 |
| Mean gene size<br>(bp) | 7226 | 6834 | 10675 | 14305 | 13329 | 13255 |
| Genes with<br>functional<br>prediction | 24657 | 24766 | 23686 | 22597 | 22453 | 20938 |

\* Number of scaffolds > 10M nt.
\*\* %A, %C, %G, %T, %N.
\*\*\* Complete single copy and duplicated.
\*\*\*\* C: Complete; S: Complete and single copy; D: Complete and duplicated; F: Fragmented; M: Missing.
Total BUSCO groups searched: 3640 Teleostei database.
Genes with functional prediction either have a functional annotation from InterProScan or name from comparisons with UniProtKB/SwissProt.

The annotation of the genome assemblies resulted in a total number of predicted genes ranging from 22124 (of these 20938 with functional predictions) in European hake to 39969 (of these 24766 with functional predictions) in polar cod, and the BUSCO gene completeness score ranged from 88.4% to 95.3%. For a full overview of all species see Table 1. Complete mitogenomes were successfully assembled for the species included and utilized for downstream phylogenetic analyses (Supplementary note 1 and Figures S2, S3, and S4 for more details on mitogenomes).

### Macro-synteny and identification of chromosomal rearrangements

To explore the macro syntenic interspecies relationship between Atlantic cod (NEAC) and the five other codfish species, we conducted gene order comparisons. The analyses revealed in general conserved intra-chromosomal macro synteny for a majority of the chromosomes, but some larger inter-chromosomal rearrangements such as fusion, fission, and translocations were detected (Figure 2A, Supplementary note 2, Figures S5, S6, and S7 for more details). The largest reorganizations seem to have occurred within the Arctic cod and polar cod (Figures 2A and 3A), where several species-specific chromosomal fusions were observed. In total, eight fused chromosomes were detected in Arctic cod with an average size of ≥ 44 Mb, whereas five fused chromosomes were detected in polar cod with an average size of ≥ 42 Mb. These fusions were found to correspond to at least two homologous chromosomes in the other codfishes (Figure 3A and Table S3), thus resulting in the reduced number of chromosomes n=15 and n=18 found in Arctic cod and polar cod, respectively (Figure 3A and Table S3). Intriguingly, none of the fusion events seem to be of common origin as none of the fused chromosomes in Arctic cod and polar cod are 1:1 homologous.

**Figure 2:**
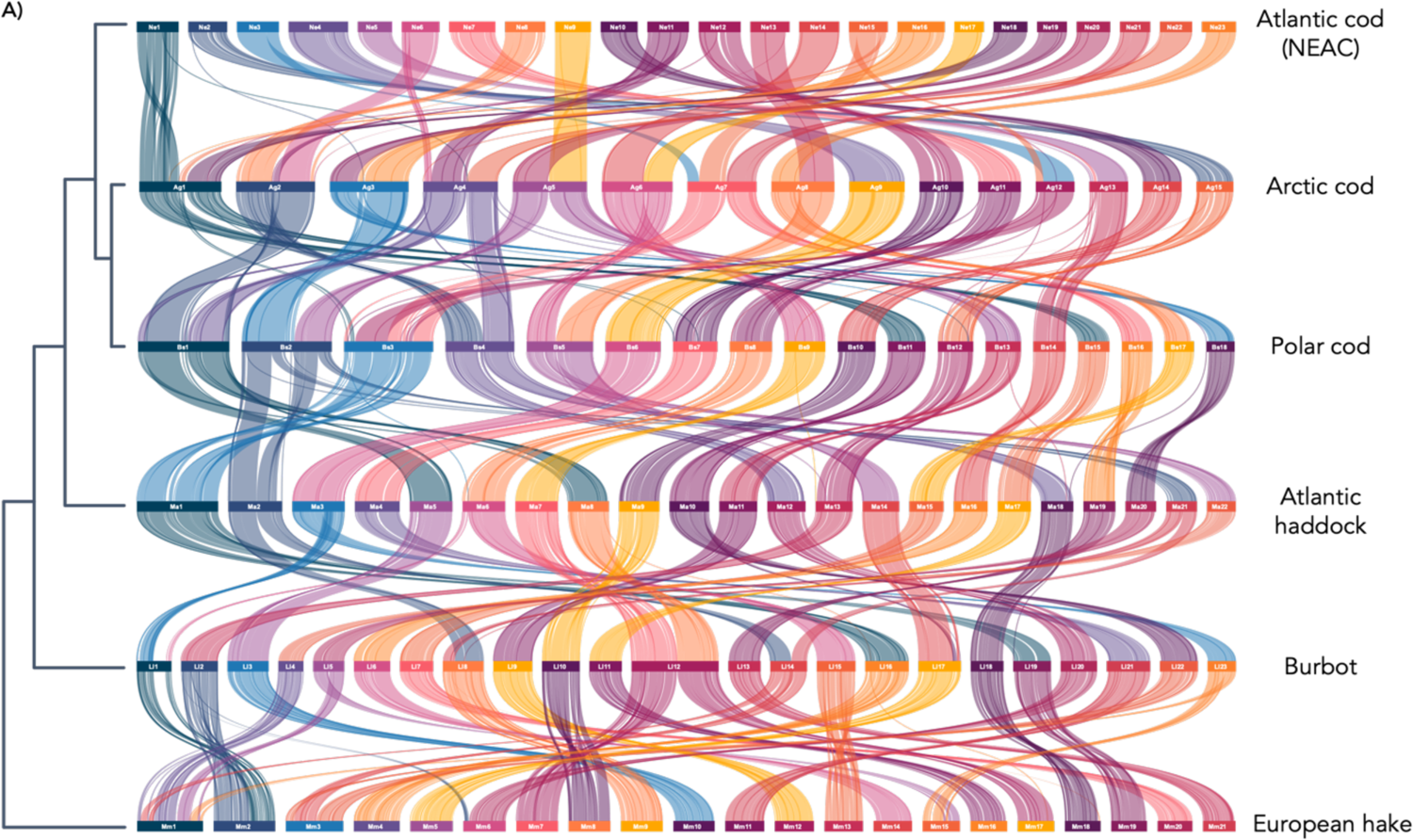
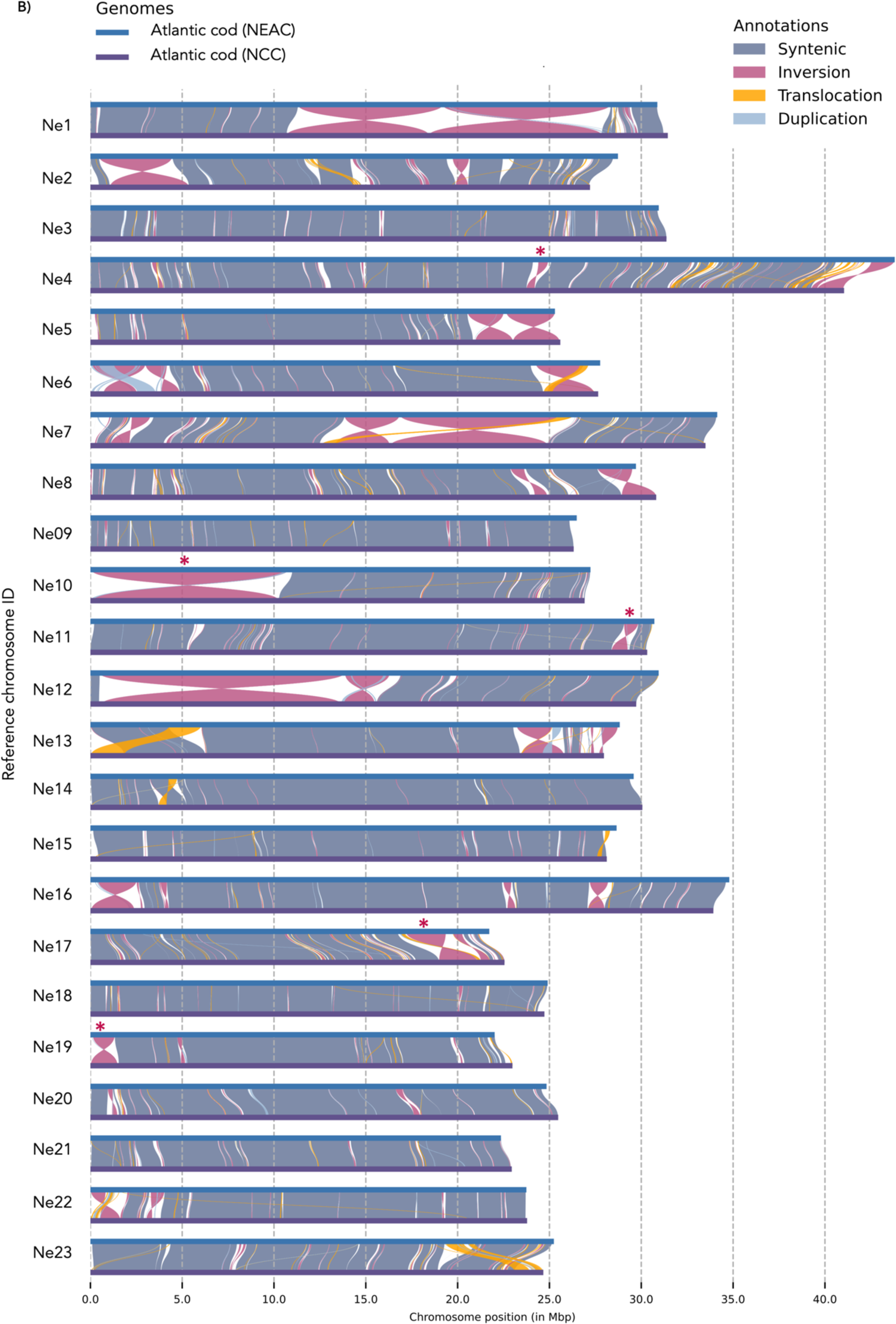
Chromosomal synteny among and within Gadiform fish species. **A)** Chromosomal synteny inferred by gene order using the programs MCScanX[116] and SynVisio[117] among the six codfishes. Chromosomes are ordered by size from largest to smallest except for Atlantic cod. Phylogenetic relationships redrawn as a cladogram based on multispecies coalescent (MSC) species tree inferred using Astral-III[138] of 1939 BUSCO gene trees shared between all taxa. **B)** Chromosomal synteny between the two Atlantic cod ecotypes the migratory Northeast Arctic cod (NEAC) and the non-migratory Norwegian coastal cod (NCC) visualized using SyRI[43]. Red stars denote putative chromosomal inversions identified in the present study, where signals of elevated linkage disequilibrium (LD) have previously been reported when populations of Atlantic cod have been examined [36,38,48].

**Figure 3:**
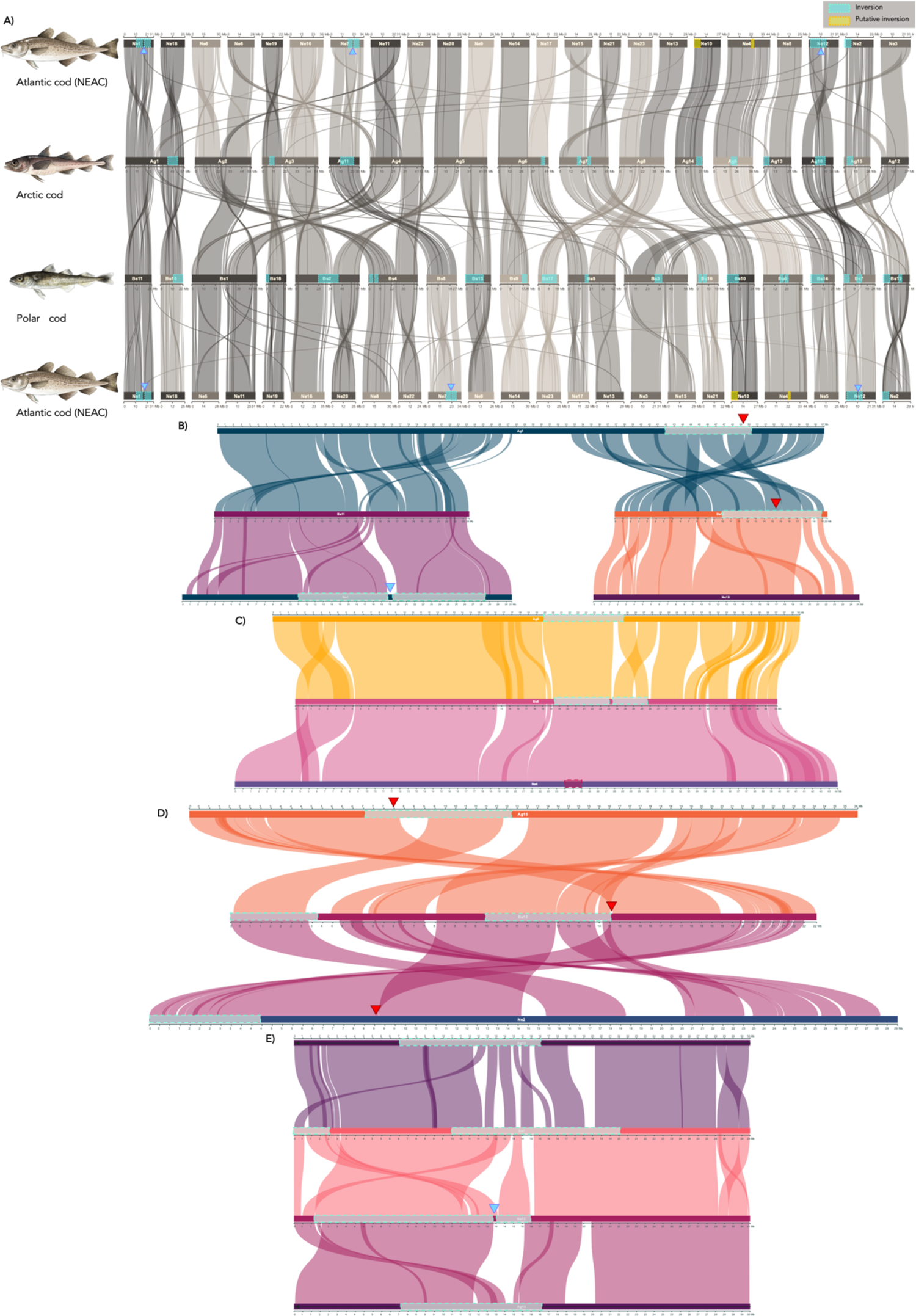
Chromosomal synteny and schematic overview of intraspecies inversions in Arctic cod, polar cod, and Atlantic cod (NEAC). **A)** Chromosomal synteny between two Arctic codfishes as well as Atlantic cod, intraspecies polymorphic inversions marked with turquoise squares, and putative inversion marked in yellow. **B-E)** Pairwise chromosomal syntenies for a selection of chromosomes where intraspecies inversions overlap between Arctic cod, polar cod, and/or Atlantic cod (NEAC). For B) Ag1 - Bs11 & Bs15 - Ne1 & Ne18, C) Ag9 - Bs6 - Ne4, D) Ag15 - Bs12 - Ne2 and E) Ag10 - Bs7 - Ne12 - Ag10. Intraspecies inversions are marked in turquoise, and one putative inversion is marked in red (on Ne4). Approximate breakpoint region between two subsequent double inversions in Atlantic cod marked with blue triangles. Locations of hemoglobin gene clusters associated with inversions are marked with red triangles, where LA and MN clusters are located in **B)** and **D)** respectively.

As mentioned above, we identified several inter-chromosomal translocations of smaller and larger genomic regions when comparing polar cod and Arctic cod vs. Atlantic cod. For instance, within the fused chromosome Ag1 in Arctic cod, which is primarily homologous to two chromosomes in Atlantic cod (Ne1 and Ne18; see Materials and Methods for the renaming of the Atlantic cod chromosomes), displayed a central region of the chromosome (∼25 – 34 Mb) which was homologous to the tail end of a third chromosome in Atlantic cod (Ne15). For the fused chromosome Ag2 in Arctic cod, which is mainly homologous to Ne8 and Ne6 in Atlantic cod, a centrally located region on Ag2 (∼22 – 30 Mb) was found to correspond to a smaller region at the end of Ne11 in Atlantic cod (Figure 3A and Table S3). Additionally, the fused chromosome, Ag4 primarily homologous to Ne11 and Ne22 in Atlantic cod, displayed a region at the beginning of the chromosome (∼1 – 5 Mb) corresponding to a chromosomal segment at the beginning of Ne6 in Atlantic cod. The fused chromosome Ag7 was found to mostly correspond to Ne15 and Ne21, whereas the first part of the chromosome (∼1 – 10 Mb) was found to be homologous to a chromosomal region in the tail end of Ne3 in Atlantic cod. For Ag12 which was primarily homologous to Ne3, a smaller region in the beginning of the chromosome (∼1 – 5 Mb) corresponds to the first part of Ne15 in Atlantic cod (Figure 3A and Table S3).

In the comparisons with polar cod, we found that the fused chromosome Bs1, mainly homologous to Ne6 and Ne11 in Atlantic cod, displayed a region centrally located on the chromosome (∼25 – 29 Mb) corresponding to the tail end of Ne8, Ne13 and the beginning of Ne22 in Atlantic cod. Finally, in the fused chromosome Bs3 in polar cod, mainly homologous to Ne3 and Ne15 in Atlantic cod, a subsequential intra-chromosomal translocation of the end region on Ne15 (∼25 – 29 Mb) to the beginning of Bs3 (∼1 – 10 Mb) has seemingly occurred (Figure 3A). However, this reshuffling could also be explained (but less likely) by an insertion of Ne3 into Ne15 with additional subsequent translocations (see Supplementary note 2 and Figures S7C and D).

Visual inspection of Hi-C heatmaps across the genome assemblies also revealed that the largest chromosomes in Arctic cod and polar cod, i.e., the fused chromosomes, centromeres appear to be centrally located for all chromosomes, except for chromosome 3 (Bs3) in polar cod (Figure S1D and E for Arctic cod and polar cod, respectively).

### Intraspecies chromosomal synteny between Atlantic cod ecotypes

The generation of the chromosome-level genome assembly for the non-migratory Norwegian coastal cod (NCC) enabled in-depth inspection of the chromosomal synteny with the already released version of the genome assembly of the migratory Northeast Arctic cod[42] (gadMor3.0, NCBI refseq assembly: GCF_902167405.1). By the comparative analyses (using SyRI[43]), we identified several smaller and medium sized chromosomal inversions (Figure 2B), in addition to the already known chromosomal inversions on LG01, LG02, LG07, and LG12[36–39,44], here synonymous with Ne1, Ne2, Ne7, and Ne12 (Figure 2B). Of the already characterized chromosomal inversions, we discover that the inversions earlier identified on Ne7 and Ne12 are both made up of double inversions (see Figure 2B). For the inversion on Ne7, a recent study has reported that single nucleotide polymorphisms within different parts of the inversion display distinctive frequencies among populations of Atlantic cod analyzed[45], possibly reflecting the double inversion identified in the present study. Furthermore, for the inversion on Ne12 discrepancies have been reported for the length of the inversion, with the latter breakpoint either being located at around 13.5 Mb[10,36,46] or alternatively around 16 Mb[39]. Here, we provide support that this inversion is a double inversion, and the length variation previously reported may reflect differences in frequency of the genotypes of the second, smaller inversion located at ∼13.5 – 16 Mb on Ne12 among the different populations examined (see Figure 2B). We provide further support for the previously identified double inversion on Ne1[47], with our analyses delineating two subsequent inverted segments (Figure 2B). Additionally, we discovered one putatively larger inversion on Ne10 (Figure 2B), not yet discovered by population genome-wide datasets. It should be mentioned, however, that within this chromosome patterns of high linkage disequilibrium (LD) have been detected, especially in comparisons where Baltic and Canadian populations were included[36,38]. For some of the smaller and medium sized inversions mentioned above, we here want to highlight the ones on Ne11, Ne17, and Ne19 (shown with red stars in Figure 2B) which are found to be overlapping with genomic regions of high LD in multiple population genomic studies[36,38,48]. Lastly, we highlight one smaller putative inversion detected on Ne4 (Figure 3A, Figure 3C, and Figure 2B), which seems to be partly overlapping with the species-specific inversions detected in polar cod and Arctic cod (Figure 3A, Figure 3C; see description below for more information). For this and some of the other inversions, population genetic analyses are needed to confirm that these identified putative inversions are polymorphic on a broader population scale.

### Genomic differentiation and detection of overlapping chromosomal inversions

Inspecting the locations of the earlier reported intraspecies chromosomal inversions in Atlantic cod[36–39,44], polar cod[34], as well as Arctic cod[35], we detected in total eight partly overlapping inversions between the two Arctic species, where two of these were also found to be partly overlapping with two of the inversions detected in Atlantic cod (Figures 3B-E and Figure S8). Further, we discovered one additional partly overlapping inversion between either Arctic cod or polar cod vs Atlantic cod (Figure 2A).

From the population data utilized in this study (i.e., Arctic cod and polar cod samples collected in Tyrolerfjorden, Greenland [N=14] and the Barents Sea [N=14], respectively) the genomic differentiation estimates and/or sequence similarity along chromosomes between the two Arctic species revealed overall high degree of differentiation as expected, with a baseline F_ST_ averaging 0.8-0.9 (see Figure 4A-D and Figure S9). However, we also identified local stretches along the chromosomes where lower values of F_ST_ (below 0.5) as well as close to zero values of genetic divergence (D_XY_) (Figure 4A-D and Figure S9) were observed. Specifically, such regions were localized in connection to the partly overlapping inversions and/or in the breakpoint regions identified (Figure 3B-E). Moreover, the estimated nucleotide diversity (π) was on average lower in Arctic cod than in polar cod (Figure 4A-D, and Figure S9), i.e., most likely a result of reference bias, since we here used Arctic cod as a reference genome for the variant calling[35].

**Figure 4:**
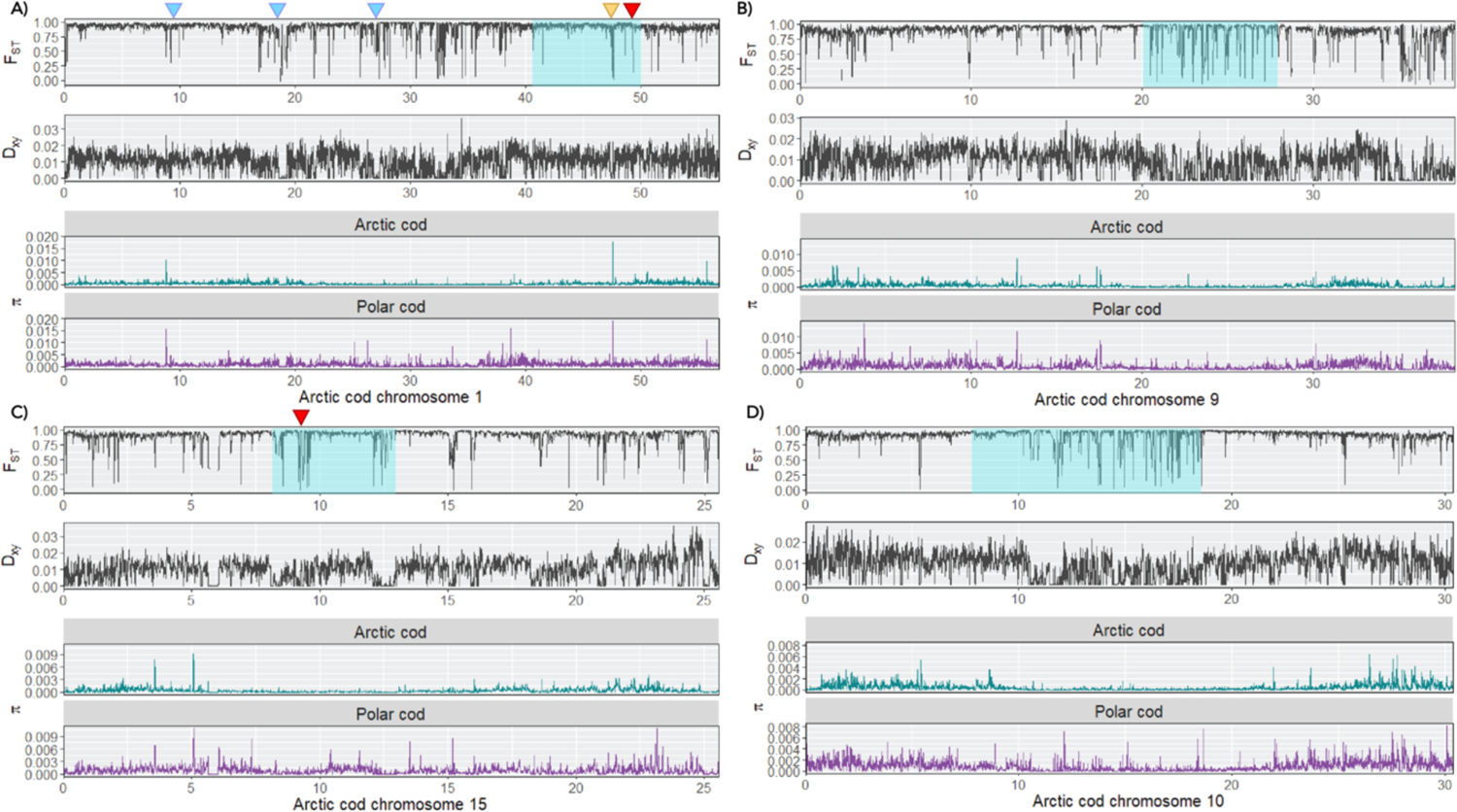
Genetic diversity and differentiation between Arctic cod and polar cod, within inversion regions. A-D) Intraspecies genetic differentiation and nucleotide diversity estimated by pixy[126] between Arctic cod and polar cod using the Arctic cod genome assembly as a reference, for chromosomes 1, 9, 15, and 10. The population data includes Arctic cod (n=14) and polar cod (n=14) collected in Tyrolerfjorden, Greenland, and the Barents Sea respectively. Blue triangles highlight approximate breakpoint regions of the double inversion in Atlantic cod on Ne1, in homologous regions of Arctic cod Ag1, yellow triangle indicate the position of FBXL5 gene (connected to homeostasis[49]), and purple triangles specify the position of hemoglobin gene cluster locations on Arctic cod Ag1 and Ag15. Turquoise squares indicate intraspecies inversions in Arctic cod.

More specifically, when comparing the last part of the fused chromosome Ag1 with the corresponding homologous chromosomes in the two other species, i.e., Bs15 in polar cod and Ne18 in Atlantic cod, we found the species-specific chromosomal inversions in polar cod and Arctic cod to be partly overlapping (Figure 3B). Intriguingly, one of the hemoglobin gene clusters (the LA cluster) was identified to be located at the very edge of the inversion in Arctic cod, while found to be located within the inversion in polar cod (see Figure 3B, where the localization of the LA cluster is defined with a blue triangle). Moreover, comparing the first part of the fused chromosome of Ag1 with the corresponding homologous chromosomes, i.e., Bs11 and Ne1, we detected no presence of any overlapping inversions between the Arctic species. For Atlantic cod, however, this is the chromosome (Ne1) where the large inversion discriminating between the non-migratory and migratory behavior is localized[10,36,38,47]. However, for the same region using Arctic cod chromosome 1 (Ag1) as the reference, by pairwise comparisons between polar cod and Arctic cod, we identified several stretches with F_ST_ estimates lower than 0.5, i.e., indicating higher degree of sequence similarity between the two species in those regions (Figure 4A). These low F_ST_ stretches were i) overlapping with the three breakpoint regions of the identified double inversion in Atlantic cod Ne1[47] (Figure 3B, Figure 4A, and Figure 2B) and ii) the breakpoint regions of the partly overlapping inversion identified between polar cod and Arctic cod (Figure 3B and Figure 4A). Inside the latter inversion, we also detected lower estimates of F_ST_ as well as high values of π for the region harboring the F-box and leucine-rich repeats protein 5 (FBXL5) gene in Arctic cod, which has been linked to iron homeostasis[49].

For the homologous chromosomes Ag9, Bs6, and Ne4 we identified species-specific inversions which overlapped between the two Arctic species as well as with a smaller putative inversion detected in Atlantic cod within this study (Figure 3C and Fig 2B). Moreover, the first breakpoint region for the overlapping inversions in polar cod and Arctic cod is homologous with a high LD region (i.e., a putative inversion) detected in Atlantic cod on Ne4[36,38,48]. Calculations of F_ST_ and D_XY_ between polar cod and Arctic cod along chromosome 9 (Ag9) of Arctic cod uncovered multiple regions with very low F_ST_ estimates (lower than 0.25) compared to surrounding regions. Here, the stretches of low F_ST_ overlap entirely with the overlapping inversions between the two Arctic species. Moreover, calculations of D_XY_ display a putatively similar pattern, with lower estimates over the same region (Figure 4B).

For the homologous chromosomes Ag15, Bs12, and Ne2 the inversion detected in Arctic cod was found to be partly overlapping with one of two chromosomal inversions detected in polar cod (Figure 3D). In addition, due to intra-chromosomal translocations, the Arctic cod inversion seems to have an overlapping breakpoint region with the second inversion detected in polar cod. Intriguingly, this genomic region enharbours the second hemoglobin cluster (the MN cluster), which is localized outside of the inversion detected in Atlantic cod, at the end breakpoint region of the inversion detected in polar cod and inside the inversion detected in Arctic cod (Figure 3D, the MN cluster is marked with a blue triangle). Moreover, estimates of F_ST_ and D_XY_ indicate higher degree of sequence similarity at the breakpoint regions of the inversion found in Arctic cod as well as for the MN cluster (Figure 4C, indicated with a blue triangle).

When comparing the homologous chromosomes Ag10, Bs7, and Ne12 we found that the species-specific inversions detected, i.e., the putative double inversion in Atlantic cod (Figure 2B) vs. one inversion in Arctic cod and two inversions in polar cod, are seemingly all overlapping (Figure 3E). In addition, intra-chromosomal translocations have further led to shared local genomic regions between the inversions, e.g., a region within the inversion on Ag10 in Arctic cod, seems to translocate and make up the first breakpoint region in the first of the two inversions identified in polar cod (Figure 3E), whereas the end breakpoint of the first inversion of the double inversion in Atlantic cod makes up the latter part of this first inversion in polar cod (see Figure 3E). Moreover, the overlapping genomic regions between the inversions in Arctic cod and polar cod, i.e., the latter part of the inversion detected in Arctic cod, exhibited low estimates of F_ST_ between the two Arctic species compared to surrounding regions (Figure 4D).

For the homologous chromosomes Ag11, Bs8, and Ne7 an inversion detected in Arctic cod was found to overlap to some extent with the double inversion in Atlantic cod, where the major differentiation seems to be linked to translocation of the breakpoint regions in Arctic cod (Figure 5A and C). For this region however, no signals of sequence similarity (F_ST_ nor D_XY_) were detected between polar cod and Arctic cod except for a smaller region co-localizing with the first breakpoint region in Arctic cod that is translocated in Atlantic cod (Figure 4A and C).

**Figure 5:**
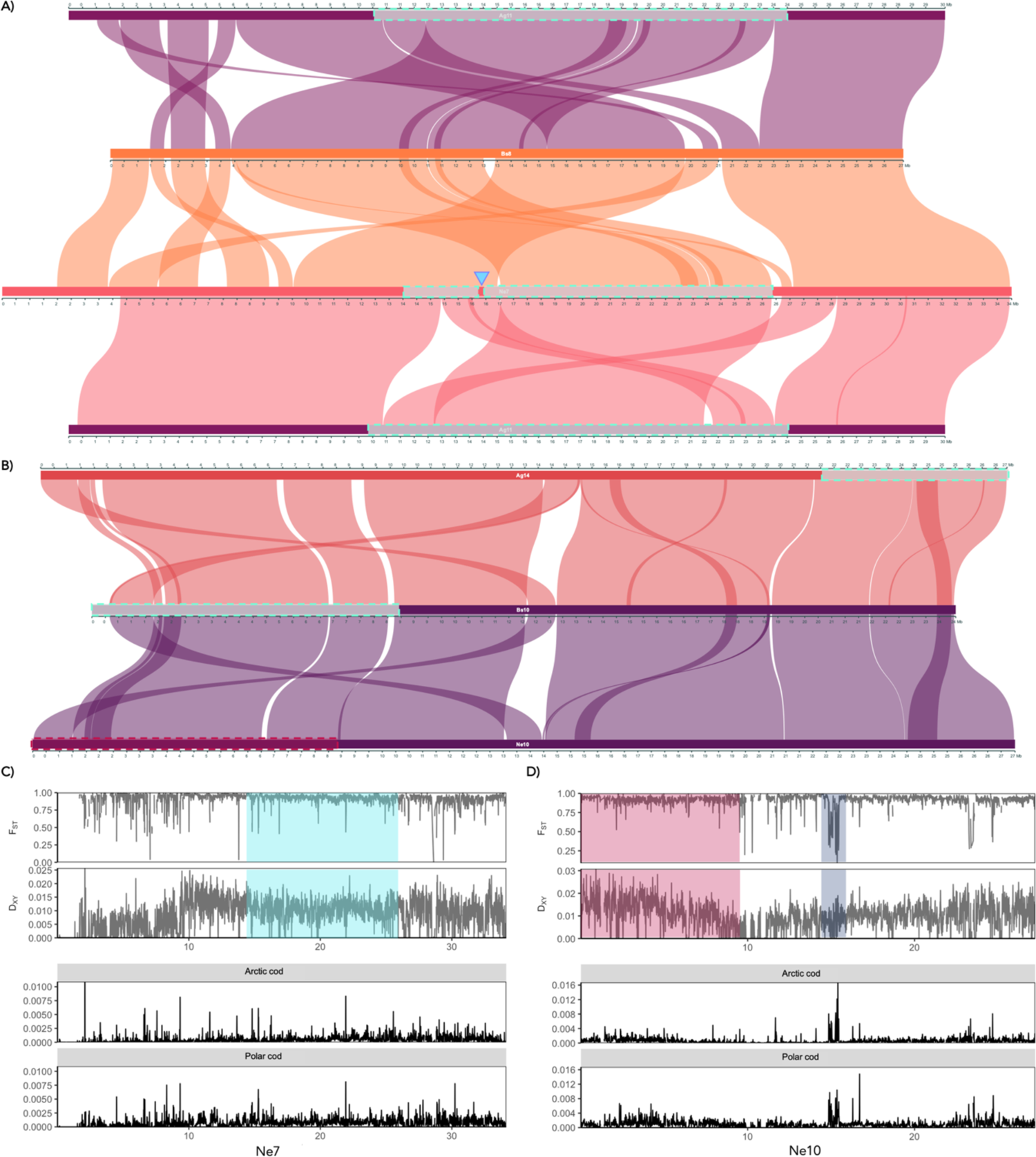
Chromosomal synteny, genetic diversity, and differentiation within inversion regions between Atlantic cod, Arctic cod, and polar cod. Chromosomal synteny between the Arctic gadids and Atlantic cod (NEAC) **A)** Ag11 - Ne7 - Bs8 - Ag11 and **B)** Ag14 - Bs10 - Ne10. The boundary between the double inversions in Atlantic cod on Ne7 is marked with a blue triangle. **C)** and **D)** Interspecies genetic differentiation and nucleotide diversity were calculated using pixy[126] between Arctic cod and polar cod using the Atlantic cod (NEAC) reference genome for Ne7 and Ne10, respectively. A region of sequence similarity between Arctic cod and polar cod corresponding to the first part of the inversion on Bs10 is highlighted in gray on Ne10. The intraspecies polymorphic inversion region in Arctic cod (largely overlapping with the inversion in Atlantic cod (NEAC) on Ne7) is visualized with turquoise square, while the putative inversion on Ne10 in Atlantic cod (NEAC) is marked in red.

The final comparison conducted between homologous chromosomes was between polar cod Bs10 and Atlantic cod Ne10. In polar cod, this chromosome harbors a polymorphic inversion that overlaps to some extent with the inversion detected when comparing the genome assembly for the non-migratory NCC with the migratory NEAC (gadMor3.0) (see Figure 5B and Figure 2B). In Arctic cod, an inversion has been detected on Ag14 (homologous to Bs10 and Ne10) at the end of the chromosome, however, this inversion was not found to overlap with the inversions found in polar cod and Atlantic cod (Figure 5B). Inspecting the estimates of F_ST_ and D_XY_ within this region between polar cod and Arctic cod, we detect lower values of F_ST_ for the stretches that are presumably overlapping with breakpoint regions of this inversion, as well as lower estimates of D_XY_ (∼9 Mb for the second breakpoint and 13 – 14 Mb for the translocated area making up the first breakpoint, Figure 5D).

### Elucidating the relationship of the Arctic codfishes

A Maximum Likelihood (ML) phylogenetic tree based on the concatenated protein-coding genes from the mitochondrial genomes of an extended number of species (see Materials and Methods for full list), placed Arctic cod as sister taxon to the Atlantic cod (*Gadus morhua*), although with low support, Shimodaira-Hasegawa-like approximate likelihood ratio (SH-alrt) of 26.6 and ultrafast bootstrap approximation (UFBoot[50]) of 56 (Figure 1). Pairwise identity analysis between *Melanogrammus aeglefinus*, *Arctogadus glacialis, Boreogadus saida* and *Gadus morhua* for each mitochondrial protein-coding genes (PCGs) revealed that the PCGs between *Arctogadus glacialis, Boreogadus saida* and *Gadus morhua* had high pairwise similarity (Figure S10), suggesting that the PCGs might not provide sufficient information for resolving the branch represented by *Arctogadus glacialis*.

The reduced ML and Bayesian phylogenies inferred using only the mitogenomes of the species sequenced in the present study (except for Atlantic haddock, see Supplementary Materials and Methods for details) and using concatenated PCGs resulted in the same topology as the extended mitochondrial phylogeny (Figure 1, S11, and S12).

Moreover, the ML and Bayesian tree inferences based on the complete mitogenomes differed in the placement of Arctic cod, where the ML analysis placed Arctic cod and polar cod as sister species with a bootstrap support of 62, whereas the Bayesian phylogeny placed Arctic cod as sister taxon to the Atlantic cod, with a posterior probability of ∼0.76 (Figures S13 and S14).

The BUSCO search uncovered between 88.4% and 93.9% complete and single-copy BUSCO genes for the genome assemblies presented in this study, including the already published NEAC genome assembly (gadMor3.0)[42] (Table S4). In total 2494 BUSCO genes were found to be shared among all species. 555 BUSCO genes were removed after quality filtering (see Materials and Methods for details and Figure S15).

Multispecies coalescent (MSC) analysis using Astral-III of the 1939 BUSCO gene trees resulted in a species tree with good bootstrap and posterior probabilities (Figures 6A and S16). In the resulting tree, Arctic cod and polar cod were placed as sister species, and the Atlantic cod was placed as the sister taxon to this group (Figure 6A). The ML phylogeny of the concatenated BUSCO genes with an alignment length of 3,364,610 bp resulted in a tree with the same topology as the species tree (Figure S17). Quartet frequencies revealed that the majority of gene trees agree on the placement of NEAC and NCC as expected (Figure 6B and Figure S18). However, the relationship of Arctic and polar cod in relative to *Gadus morhua* show some level of discordance (Figure 6C and Figure S18) with the highest frequency of gene trees placing Arctic cod (*Arctogadus glacialis)* and polar cod (*Boreogadus saida*) as sister species (0.495), Arctic cod (*Arctogadus glacialis)* as the sister lineage to *Gadus morhua* (0.272), and lastly polar cod (*Boreogadus saida*) as the sister lineage to *Gadus morhua* (0.233). Due to the discordance, we further investigated the positioning of the BUSCO genes, which could potentially contribute to uncertainties in the phylogenetic placement of the species. We found that 148 and 314 of a total of 1939 BUSCOs were localized within the species-specific inversions in Arctic cod and polar cod, respectively.

**Figure 6:**
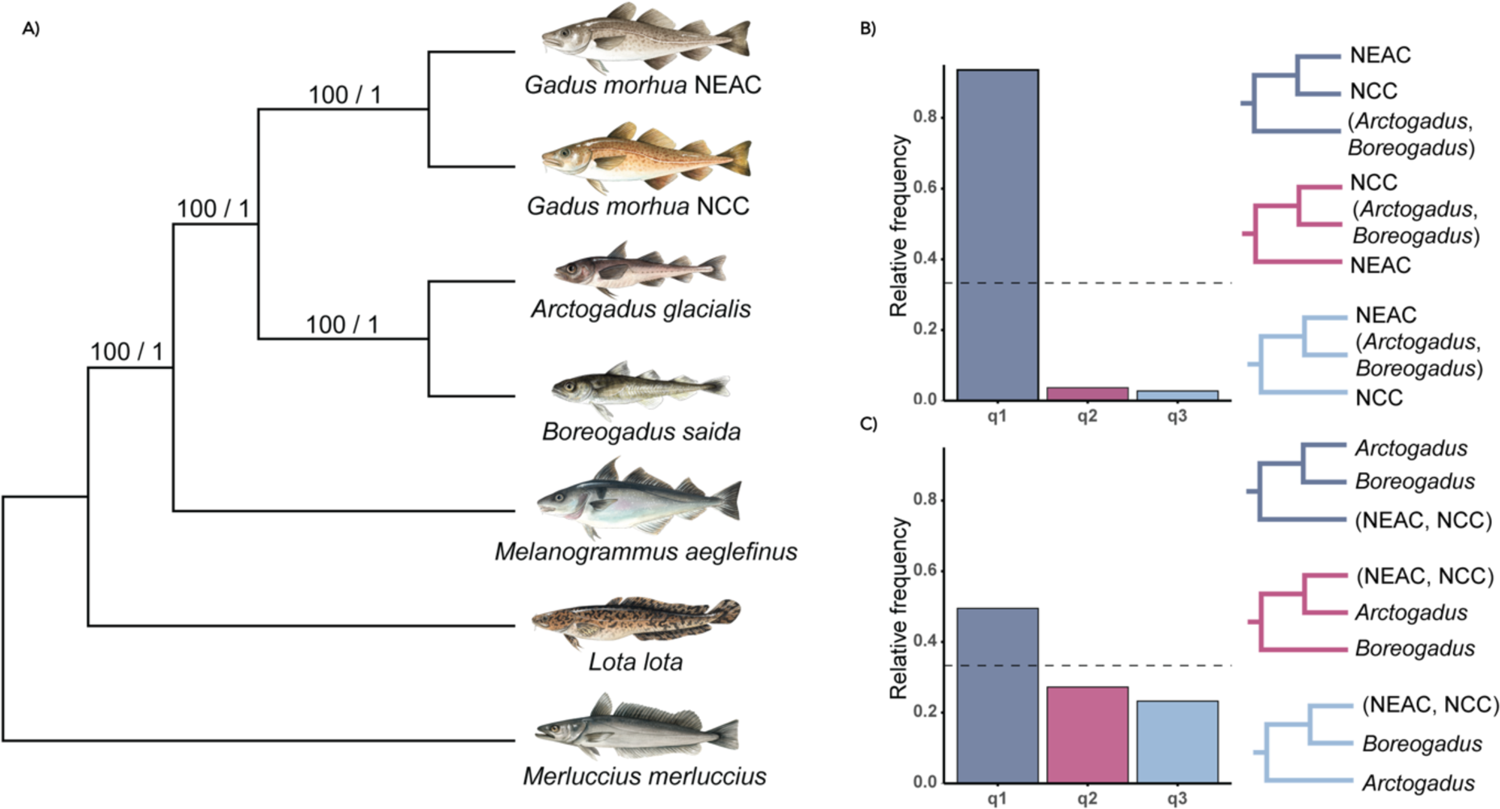
Multigene phylogenies of the Gadiformes species A) Multispecies coalescent species tree produced with Astral-III[138] of 1939 BUSCO gene trees shared between all taxa. Branch support is given as multi-locus bootstrap / local posterior probability (see Materials and Methods), and branch lengths are given as coalescent units. **B)** Quartet frequencies calculated using DiscoVista[139] with *Merluccius merluccius*, *Lota lota*, and *Melanogrammus aeglefinus* specified as outgroups. B) Quartet frequencies of the internal branch separating NEAC and NCC from the other species, and **C)** quartet frequencies of the internal branch separating *Arctogadus glacialis* and *Boreogadus saida* from the rest of the species. Each tree depicted for B) and C) represents alternative hypotheses for the unrooted quartets shown in the bar plot, outgroup not shown.

## Discussion

In this study, we have generated chromosome-level reference genomes for six codfish species to investigate the evolutionary genomic changes that have taken place within this lineage. Using a comparative framework — focusing on the Arctic gadids — we uncovered independent lineage-specific chromosomal fusions and genomic rearrangements in these species. Our results indicate that these genomic rearrangements are of evolutionary importance for these cold-water specialists.

### Chromosome number variability in codfishes

The sequencing and assemblage of reference genome assemblies for six codfish species (Atlantic cod (NCC), polar cod, Arctic cod, Atlantic haddock, burbot, and European hake) resulted in assemblies of high contiguity and BUSCO completeness metrics. The chromosome-level genome assemblies generated, corresponded to the numbers of chromosomes reported from cytogenetic studies (see Table S1), except for burbot where we obtained a total of n=23 haploid chromosomes vs. n=24 haploid chromosomes reported by two different cytogenetic studies[40,41]. We cannot rule out that super-scaffolds have erroneously been connected which should have been split in the burbot assembly, but our Hi-C contact map did not indicate any super-scaffolds that should have been further split.

Moreover, two different chromosome-level burbot assemblies have previously been constructed, which resulted in different numbers of chromosomes, one by Han et al. 2021 of a specimen sampled in Heilong River (Raohe county) in the northeastern part of China resulting in n=22 chromosomes[51], and another by Song et al. 2021 resulting in n=24 chromosomes[52]. Combined, these results suggest that there are chromosomal differences between different lineages of burbot. Previous studies using mitochondrial markers point towards at least five major mitochondrial burbot clades across western and northern Europe, Russia, and parts of central Asia, likely reflecting postglacial colonization and the geological history of burbot in separate Holarctic regions[53]. Based on this, we speculate that the differences in chromosomal numbers observed might be linked to true karyotype differences and thus, different postglacial colonization events.

Cytogenetic studies of Gadiform species have showcased an extensive karyotype variability among species in the group (Figure 1 and Table S1), with chromosomal numbers ranging from n=13 to n=24. Our analyses revealed that independent chromosomal fusion events have taken place within the lineages of Arctic gadids, encompassing the largest five vs. eight chromosomes within polar cod and Arctic cod, respectively. It is rather striking that none of the fusion events within each of the lineages seemed to be of common origin, as none of the fused chromosomes were homologous between the species. A reduced set of chromosomes has also been reported in two codfish species residing in Arctic and sub-arctic waters[25,54,55], i.e., the Navaga (*Eleginus nawaga*) and Saffron cod (*Eleginus gracilis*) (Figure 1 and Table S1). Another lineage that displays high degree of karyotype variability among species, and thus, a pronounced reduction in numbers of chromosomes in some lineages, are the cold-water adapted Antarctic notothenioids[21–23,56]. Taken together, the reduction in number of chromosomes suggests an adaptive significance of chromosomal fusions in species inhabiting high-latitude oceans, thus could be examples of convergent/parallel evolution in response to cold adaptation.

Using the Hi-C contact map we were able to localize centromeres for most of the fused chromosomes to the center of the fused regions (see Figure S4D and E), i.e., being metacentric. This was the case for all except chromosome 3 (Bs3) in polar cod, where the centromere was localized at one end of the chromosome possibly due to a translocation, i.e., being acrocentric. Thus, the majority of fused regions in the Arctic gadids co-localize with the centromeres, which are regions suggested to display reduced recombination rates (and therefore exhibit elevated LD) compared to surrounding regions across many taxa[57,58] (reviewed in Choo[59]).

A population sequencing study of polar cod demonstrated that the five chromosomal fusions reported here, display unique genomic characteristics, including signals of elevated LD, positive estimates of Tajima’s D, and reduced D_XY_ estimates compared to surrounding regions[34]. Similar results were also seen, but to a lesser extent, in a small-scale population study of Arctic cod[35]. The signals of higher LD indicate lower recombination rates within these regions at the center of the chromosomes vs. lower LD signals and thus, higher recombination rates are detected further away towards the distal regions of the chromosomes. These findings are in line with previous studies reporting that chromosome size (due to fusions) and localization of the new centromere, impact the recombination landscape across chromosomes[60–62]. Reduced recombination across a fused region of a chromosome, is likely to have evolutionary consequences, i.e., by increasing the overall linkage between genes and genetic elements residing within this region, i.e., leading to genes being inherited together more frequently. Chromosomal fusions have the ability to bring together previously distantly located genes and genetic elements that may be of particular advantage in a given environment[63]. Therefore, rearrangements of chromosomes may give rise to new and advantageous combinations of genes/genetic elements, as well as increase the likelihood of the new combinations being inherited together. In this regard, we highlight that three of these fused regions in polar cod harbor inversions, i.e., on Bs2, Bs3, and Bs5, and for the latter the very same region has also been identified to harbor a putative sex-determining region in polar cod[34], further supporting that these regions contain genes of potentially evolutionary importance.

Moreover, direct beneficial roles of chromosomal fusions themselves, due to the increased probability of co-adapted alleles being inherited together as a result of chromosome fusions and thus, lowering the risks for segregation errors as seen with higher chromosome numbers[62], should not be neglected. Especially within the Arctic gadids, where multiple larger species-specific chromosomal inversions are identified[34,35], and the survival rate in general is low due to inhabiting highly variable and harsh conditions[64], such segregation errors could be lethal. Lastly, it should be mentioned that the unique chromosomal fusions detected for the partially sympatric sub-Arctic and Arctic gadids[25] can act as barriers to gene flow (speciation by chromosomal rearrangements discussed in[63,65–67]), and thus, be linked to the diversification and speciation processes within the lineage.

### Chromosomal inversions in the Arctic gadids

The in-depth investigation of the genomic localization of the previously reported intraspecific chromosomal inversions found in Atlantic cod, polar cod, and Arctic cod uncovered multiple partly overlapping inversions among the species, as well as for some of the breakpoint regions (Figure 3A for overview, and B-E for detailed examples). None of these inversions was 100% overlapping, i.e., indicating that the chromosomal inversions detected are species-specific, and thus of evolutionary independent origin. However, given the fact that local genomic reorganizations seem to have happened regularly within the different species, e.g., such as inter- and or intra-chromosomal translocations, a common origin for some of the inversions cannot be ruled out completely. For instance, the rather large inversion detected when comparing the genome assemblies of the non-migratory and migratory Atlantic cod and the corresponding inversion detected in polar cod on chromosome 10 (Ne10 vs. Bs10) could potentially have a shared ancestral origin, followed by successive translocations within polar cod (Figure 5B). Similarly, a common origin cannot be excluded for the partly overlapping inversion found between polar cod and Arctic cod on Bs6 vs. Ag9, as well as for the inversions found to be overlapping to some extent between Arctic cod and Atlantic cod on Ag11 vs. Ne7. For the latter, the discrepancy is by large due to the translocation of the breakpoint regions within Arctic cod. Limited sequence similarities (F_ST_ and D_XY_) were detected over the inversion regions for the ones identified in either polar cod or Arctic cod vs. Atlantic cod (on Ne10 and Ne7). This is likely due to polar cod and Arctic cod not sharing overlapping inversions in the specific regions investigated. For the rest of the partly overlapping inversions, we uncovered smaller and/or larger local regions with high sequence similarity, indicating that the inversions harbor genomic regions with high degree of conservation between the species. Such conservation implies that these genomic regions likely harbor genes or regulatory elements of evolutionary importance that are either under strong i) purifying selection, ii) parallel selection in polar cod and Arctic cod[68,69], and/or iii) due to past gene conversion events[10]. Since polar cod and Arctic cod have overlapping geographical distributions[64,70], it is plausible to assume that they encounter comparable environmental conditions, and thus similar selection pressures acting on the same genomic regions. Some of these conserved regions span almost the entire inversion region, whereas others include smaller regions often located at the breakpoint regions of the inversion. Our results suggest that the genes found in these regions are of high evolutionary importance, either i) selected for as a supergene (and thus suppressing recombination in the inverted region) encompassing locally adapted genes/alleles, or ii) that important genes are located at the breakpoint regions — and with the formation of an inversion — advantageous gene variants are linked together[71].

Moreover, we uncover that several of the inversions, even if not completely overlapping, often harbor homologous breakpoint regions frequently in combination with translocations. Our findings suggest that these translocated syntenic regions include genomic structures that are likely to be involved in the formation of chromosomal inversions, such as transposable elements[72–75]. Codfishes are known to harbor large amounts of repeated sequences, in particular short tandem repeats[76–79] as well as transposable elements[79]. It is possible that the shared homologous regions underlying chromosomal inversions contain such structures and are mechanistic in repeated independent formation of inversions in the different species. Manual investigation of the localization of the previously described hemoglobin gene clusters, the LA- and MN-clusters[80], revealed that both were found in association with the partly overlapping species-specific inversions in the Arctic gadids, and give further support for the adaptive significance of the genomic reorganizations within codfishes. The hemoglobin genes have been linked to temperature adaptation through polymorphisms and copy-number variation in codfishes[81–84]. Interestingly, the LA-cluster was localized at the breakpoint region of the inversion on Ag1 in Arctic cod, while the cluster was found inside the inversion in polar cod (Bs15). One hypothesis for the maintenance of polymorphic chromosomal inversions after the inversion occurs states that inversions can rise in frequency within populations due to beneficial mutations close to or at inversion breakpoints[71]. Moreover, genes proximal to inversion breakpoints may exhibit differential expression[85], and genes located at breakpoints can contain different exon sequences in the alternative inversion haplotypes[86]. Based on this, we can hypothesize that hemoglobin and co-regulated genes (such as the FBXL5 gene) could be affected by the inversion(s) on Ag1 and Bs15, but to what extent remains to be explored.

The second hemoglobin cluster, the MN-cluster, was found to be localized inside the inversion on Ag15 in Arctic cod while on the boundary/breakpoint of one of the two inversions located on Bs12 in polar cod. These chromosomes are homologous to Ne2 of Atlantic cod, where an inversion also has been detected[36,38,44] — even if localized together with the MN-cluster — there is seemingly a link between this inversion and the hemoglobin genes also in Atlantic cod[87]. Moreover, the inversion on Ne2 on Atlantic cod has been linked to adaptation to environmental differences in oxygen, salinity, and temperature between Atlantic cod populations in the North Sea and the Baltic Sea[48,88]. Taken together, the MN-cluster was found to be in close association with species-specific inversions in all three species, which suggests that these homologous regions are likely of evolutionary importance in the different species. However, further investigation is needed to fully resolve the evolutionary significance of these inversions and how they relate to hemoglobin functionality.

### Are phylogenetic investigations hampered by the chromosomal rearrangements?

The phylogenies constructed using BUSCO as well as mitochondrial genes extracted from the newly generated genome assemblies, uncovered some uncertainty in the placement of the Arctic gadids. Depending on the markers and methodology used, Arctic cod was either placed as sister taxon to *Gadus*, or as sister species to polar cod, consistent with incongruences earlier reported[10,89–91]. At the mitochondrial level, our results show that the Arctic cod is a sister taxon to *Gadus*, but with low bootstrap support. The low support may be attributed to purifying selection on mitochondrial PCG (shown by Wilson et al.[91]), as the mitochondrial PCGs’ pairwise identities were found to be very similar in Arctic cod, polar cod, and Atlantic cod (Figure S10). Conversely, in the MSC BUSCO phylogeny, Arctic cod is placed as a sister species to polar cod. The quartet frequency analysis for the BUSCO phylogeny demonstrated discordance among the gene trees in the placement of Arctic cod and polar cod as sister lineage *Gadus* (Figure 6C). The uncertainty in the placement of Arctic cod in the BUSCO phylogeny could be due to several factors affecting phylogenetic inference, e.g., introgressive hybridization[10,92], incomplete lineage sorting, and/or large amounts of highly linked genomic regions leading to an altered recombination landscape[93]. For the latter, quantification of BUSCOs within and outside identified chromosomal inversions in polar cod and Arctic cod revealed some BUSCOs to be localized within these regions, possibly contributing to uncertainties in the phylogenetic placement of the species.

## Conclusions

In this undertaking, we uncover that two keystone Arctic codfishes, polar cod, and Arctic cod, have undergone extensive genomic rearrangements, including chromosomal fusions as well as inversions, likely to have played important roles in the speciation processes within the lineage, and/or adaptation to freezing environmental conditions. Finally, our attempt to reconstruct the phylogenetic relationship between the codfishes resulted in conflicting topologies for the BUSCO and mitochondrial phylogenies. We speculate that incongruences in the placement of Arctic cod might, at least partly, stem from the massive genomic reorganizations that have taken place, which cover large proportions of the genomes, encompassing a combination of shared and unique evolutionary trajectories.

## Materials and Methods

### Sample collection

We whole genome sequenced a total of six codfish species in the present study. These specimens, along with their sampling locations, are as follows: Arctic cod (*Arctogadus glacialis*) from Tyrolerfjorden, Northeast Greenland; polar cod (*Boreogadus saida*) from Tyrolerfjorden, Northeast Greenland; non-migratory Atlantic cod (*Gadus morhua*) i.e Norwegian coastal cod (NCC) from Lofoten area, Norway; Atlantic haddock *(Melanogrammus aeglefinus)* from Tromsø area, Norway; burbot (*Lota lota*) from Lake Stuorajavri, Finnmark, Norway; and European hake (*Merluccius merluccius*) from Oslofjorden, Norway. For a detailed overview of the specimens, as well as the tissue used for each sequencing technology see Table S5.

### Generation of chromosome-anchored genome assemblies of six codfish species

To construct high-quality chromosome length genome assemblies, we leveraged a combination of PacBio long-read sequencing, and chromosome conformation Hi-C reads to scaffold contigs (see Supplementary Materials and Methods for detailed description of sequencing and assemblage). For some of the species, we in addition utilized 10X linked reads to link contigs, and Illumina sequencing data for polishing, to ensure high base quality. In brief, first, primary assemblies were constructed from PacBio long reads using Flye[94]. Next, for Arctic cod, polar cod, and Atlantic cod (NCC), to assemble primary contigs into chromosome length scaffolds, contigs were first scaffolded using 10X linked reads and the program Scaff10X[95]. For all species assemblies, Hi-C linked reads were mapped and processed using either 3D-DNA[96] or YaHS[97] (Atlantic haddock). The 3D-DNA/YaHS draft assemblies were visualized and manually inspected in the Juicebox program suite[98] or PretextView[99] (Atlantic haddock). Manual splitting of scaffolds that were assembled, but that exhibited low levels of Hi-C contact points between them was performed. Lastly, for Arctic cod, polar cod, burbot, Atlantic haddock, and Atlantic cod (NCC), the assemblies were polished using both long and short-read data. For long-read polishing, pbmm2[100] was used to map PacBio reads, and polishing was done using gcpp[100]. Lastly, Illumina paired-end reads were mapped by using minimap2[101], and subsequent base calling and polishing were done using Freebayes[102]. Finalized assembly metrics were calculated using the assemblathon_stats script[103], and completeness was assessed by the Benchmarking Universal Single-Copy Orthologs (BUSCO) software[104]. Moreover, the full mitochondrial genomes of the six Gadiform species were assembled using either MitoVGP[105] or MitoHiFi[106], depending on the available input data (see Supplementary Materials and Methods).

The reference genome assemblies were annotated using either i) a combination of funannotate[107] with RNA sequencing data processed with HISAT v2.1.0[108], Portcullis v1.2.0[109], and Mikado v2.0rc6[110], or ii) a combination of miniprot v0.5[111], funannotate v1.8.13[107], EvidenceModeler v1.1.1[112], AGAT[113] and InterProScan v5.47-82[114]. For more details see Supplementary Material and Methods.

### Genome-wide synteny analyses among codfish genomes

Based on previous studies, the suggested ancestral number of chromosomes in teleosts has been estimated to be n=24-26[31–33]. Based on this, we conducted a first screening to evaluate the chromosome configuration of Atlantic cod (n=23) by dot plotting the Northeast Arctic cod genome assembly[42] (gadMor3.0, NCBI refseq assembly: GCF_902167405.1) against platyfish (n=24) (*Xiphophorus maculatus,* GCF_002775205.1_X_maculatus-5.0, NCBI refseq assembly: GCF_002775205.1) as well as John Dory (n=22) (*Zeus faber,* GCA_960531495.1_fZeuFab8.1, NCBI refseq assembly: GCA_960531495.1) using minimap2 v2.24[101] and the D-genies interactive platform[115]. This initial screening confirmed that the chromosome configuration in Atlantic cod is similar to the outgroup species, with only a few putative fusions, encompassing two of the Atlantic cod chromosomes (for more details see Supplementary note 2).

Intraspecies chromosomal synteny between Atlantic cod (NCC) and (NEAC) was investigated by first obtaining an alignment file between the genome assemblies using Minimap2 v2.17[101] with settings -ax asm5 –eqx, then chromosomal rearrangements were identified and visualized using the software SyRI v1.5[43].

Subsequently, by a gene order based approach using the program MCScanX[116] and visualization using the SynVisio[117] online platform, chromosomal synteny between the genome assemblies of the six codfishes was performed. Here, genome collinearity, chromosomal homologies, and chromosomal rearrangements were manually inspected and identified in a pairwise manner with Atlantic cod, polar cod, and Arctic cod as references, and the five other species as queries.

The MCScanX analysis was performed according to developers’ recommendations, first doing reciprocal all by all BLAST[118] search between all proteins of the annotated gene for all the six species, using BLAST+ v2.11.0[118] with the settings: blastp -e 1e-10 -b 5 -v 5 -m 8. The hits were afterward anchored to a bed-like file made from the annotation GFF file, with information about gene names and the genomic locations of each gene. Synteny blocks were then built using MCScanX with default settings i.e., gene gap size set to 10. Results were visualized on the SynVisio interactive webpage as syntenic maps between the genome assemblies.

### Detection of overlapping chromosomal inversions

From previous literature and ongoing studies, several chromosomal inversions have been detected within Atlantic cod[36–39,44], polar cod[34], and Arctic cod[35]. For Atlantic cod, four major inversions have been detected: on LG01, LG02, LG07, and LG12[36–39,44] (synonymous with Ne1, Ne2, Ne7, and Ne12 in the present study), where the inversion on Ne1 has been linked to differentiation between the migratory Northeast Arctic cod vs. the non-migratory Norwegian coastal cod. In recent studies, a total of 20 inversions in polar cod[34] and 11 inversions in Arctic cod[35] have been reported and suggested linked to cryptic ecotypes and/or local adaptation.

Using this information, and plotting the location for the different inversions by position within the genome assembly of each of the species, they were identified onto their respective chromosomes using the SynVisio[117] interactive homepage (as described above). We then inferred if some of these inversions were overlapping or not between these three species (see Figure 3A). For visualization as well as simplicity, the chromosome numbering of Atlantic cod is renamed from linkage groups LG01-LG23 to chromosome numbers Ne1-Ne23 in this paper (see Figures 2, 3, and 5).

### Genomic differentiations along the chromosomes (F_ST_, D_XY,_ and π) for the Arctic gadids

To evaluate genetic divergence and/or nucleotide similarity along chromosomes between Arctic cod and polar cod, we took advantage of population-level data generated in Maurstad et al.[35] and Hoff et al.[34]. A total of 14 Arctic cod and 14 polar cod samples (Table S6) were selected for a joint variant calling using Arctic cod as the reference, as well as Atlantic cod[42] (gadMor3.0, NCBI refseq assembly: GCF_902167405.1). First, Illumina pair-end reads were trimmed using Trimmomatic v0.39[119] using default settings. Mapping was done using the Burrows-Wheeler Alignment Tool v0.7.17[120] (BWA-MEM algorithm) with default settings. Alignment files for each sample were merged and sorted using SAMtools v1.9[121]. Reads of duplicated origin were marked using MarkDuplicates v2.22.1[122]. Variant calling and hard filtering of variant sites was performed using the Genome Analysis Toolkit (GATK) v4.2.0.0[123,124]. First, each mapped sample was individually called into GVCFs using the “HaplotypeCaller” tool. Second, GVCFs were imported into a GenomicsDataBase using the “GenomicsDBImport” tool. Joint genotyping was performed using the GenotypeGVCFs tool to produce final VCFs. Single nucleotide polymorphisms (SNPs) were extracted and down-sampled to 100,000 SNPs using “SelectVariants” to generate diagnostic plots for specific filter parameter evaluation. Filtering was done by following the GATK hard-filtering recommendations as well as by manually inspection of the the diagnostic plots as suggested in Danecek et al.[125]. After the initial round of filtering, VCFs were further filtered using VCFtools v0.1.16[125]. Using the resulting SNP variant dataset, we calculated F_ST_, D_XY_ between polar cod and Arctic cod, as well as π for each species using pixy v1.2.6[126] [127] with a window size of 10,000 bp.

### Phylogenomic placement of Arctic cod

#### Mitochondrial phylogenetics

For our mitochondrial phylogenetic analyses, we included mitochondrial genomes for additional gadid species gathered from NCBI (see Table 1 for an overview) in addition to the assembled mitochondrial genomes for Arctic cod, polar cod, Atlantic cod (NEAC and NCC), Atlantic haddock, burbot, and European hake. Mitochondrial protein-coding genes (PCGs) were identified by MitoFish[128–130], aligned using MAFFT v7.453[127], and manually inspected and corrected for reading frame shifts before they were concatenated with PhyKIT v1.11.7 create_concat[131] to produce a supermatrix. A maximum likelihood (ML) tree was inferred using IQ-Tree2 v2.2.0[132] and ModelFinder Plus (MFP)[133] was used to search for the best substitution model under the Bayesian information criterion (BIC). Branch supports were calculated using 1,000 replicates of ultrafast bootstrap approximation (UFBoot)[134]. Pairwise similarity for each PCG was calculated among *Arctogadus glacialis*, *Boreogadus saida,* and the species included from *Gadus* using PhyKIT v1.11.7[131] (Figure S10). In addition, we conducted phylogenetic analysis including only our assembled mitogenomes (see Supplementary Materials and Methods for details).

#### BUSCO phylogeny

Single-copy orthologous genes from five of the genome assemblies (Arctic cod, polar cod, Atlantic cod (NCC), burbot, and European hake) in addition to Atlantic cod [42] (gadMor3.0, NCBI refseq assembly: GCF_902167405.1) and Atlantic haddock (ENA: ERR1473879) were identified using BUSCO v5.0.0[104]. BUSCO was run using AUGUSTUS v3.2[135] applying the lineage dataset Actinopterygii (actinopterygii_odb10) consisting of 3640 BUSCO groups. All nucleotide BUSCO genes found as single-copy and occurring within all the assemblies were extracted and aligned using MAFFT v7.453[127] with the L-INS-i strategy. Each alignment was trimmed using the smart-gap approach with ClipKIT v1.3.0[136]. Filtering of alignments included filtering on sequence length, variable sites (VAR sites), and parsimony informative sites (PI sites), as these properties have been shown to affect phylogenetic signals[137]. Sequence statistics were calculated using PhyKIT v1.11.7[131]. Filtering on sequence length (> 500 bp), VAR sites (>5%), PI sites (>2.5%), and percentage of gap sites (<=30%) removed 555 BUSCO genes (Figure S15). Alignments that were above 500 bp, contained more than 5% VAR sites, 2.5% PI sites, and had more than 70% sites without gaps were included for downstream analysis.

Phylogenetic placement of Arctic cod using BUSCO genes was inferred under both concatenation and gene tree approaches. Gene trees were first inferred using IQ-Tree2 v2.2.0[132] under the GTR+I+G substitution model[132]. Tree support for each gene was calculated using 1,000 replicates of UFBoot approximation. Species tree estimation based on the gene trees produced in IQ-Tree2 was inferred under an MSC analysis conducted in ASTRAL-III v5.7.8[138]. Bootstrap support was estimated using 100 replicates of multi-locus bootstrap based on the bootstrap trees produced with IQ-Tree2. Discordance among BUSCO gene trees was visualized using quartet frequencies with DiscoVista[139] while specifying European hake, burbot, and Atlantic haddock as outgroups.

The same BUSCO genes used to infer the species tree were concatenated into a supermatrix using PhyKIT v1.11.7[131], and phylogenetic tree inference was run in IQ-Tree2 v2.2.0 under the same GTR+I+G substation model with 1,000 replicates of UFboot.

## Author contributions

SJ and SNKH conceptualized the study. SNKH, MFM, and OKT handled, processed, and analyzed the data: SNKH generated and finalized genome assemblage for polar cod, Arctic cod, Atlantic cod (NCC) and burbot, conducted all chromosomal synteny analyses (interspecies, intraspecies and outgroup comparisons); MFM generated the European hake genome assembly, mitogenomes, performed phylogenetic analyses (BUSCO and mitochondrial) and population genetic diversity analyses; OKT performed genome assemblage (polar cod, Arctic cod, Atlantic cod (NCC) and burbot), generated the genome assembly of Atlantic haddock, and conducted the annotation for all genome assemblies. KP (Arctic cod, polar cod, and burbot) and PRB (Atlantic haddock) provided specimens for genome sequencing. SJ was in charge of sampling of the European hake and Atlantic cod (NCC). Funding acquisition by SJ, KP, and KSJ. Visualization and design of figures by SNKH, MFM, and SJ. SNKH and SJ wrote the original manuscript, with relevant sections contributed by MFM and OKT. All co-authors read, provided feedback, and improved the manuscript.

## Supporting information

Supp_comparative_genomics_of_codfishes

## Acknowledgments

Library preparations and sequencing were performed by the Norwegian Sequencing Centre, Oslo. The computations were performed on resources provided by Sigma2 - the National Infrastructure for High Performance Computing and Data Storage in Norway. We thank Alexandra Viertler for the codfish illustrations.

