## Supplementary material for "Chromosomal fusions and large-scale inversions are key features for adaptation in Arctic codfish species": Supp_comparative_genomics_of_codfishes

#### Supplementary note 1

##### Long-read mitogenome assemblies for six codfishes

Complete mitogenomes were assembled for six of the species included (Figure S2), using either MitoVGP[1] or MitoHiFi[2] (see Supplementary Materials and Methods). The European hake was an exception as the mitogenome was found to be fully assembled during the Flye assemblage stage, 16,968 bp in size, as would be expected from a complete mitogenome. Upon inspection, read depth distribution among the assemblies differed markedly (Figure S3), however, this is expected as HiFi reads (for European hake and Atlantic cod (NEAC)) are generated from collapsing sequencing reads, leaving fewer but highly accurate reads. MitoFish annotation of the mitogenome assemblies resulted in the identification of 13 mitochondrial protein-coding genes (PCGs) within all assemblies (Figure S2). In Arctic cod, the T-P spacer found before the D-Loop has been shown to contain duplicated repeat motifs[3]. The length of this region is known to differ within individuals[3]. Interestingly, the Arctic cod mitogenome assembly was found to include a stretch of repeated sequence located between tRNA-Thr and tRNA-Pro (T-P spacer) and the sequence reads within this region varied with several large indels, an indication that this area could potentially represent mitochondrial heteroplasmy (Figure S4).

#### Supplementary note 2

##### Macro-synteny and identification of chromosomal rearrangements

While evaluating chromosomal architecture between Atlantic cod (NEAC)[4] and two species outgroups (platyfish *[Xiphophorus maculatus]* and John Dory *[Zeus Faber]*), we found a few larger inter-chromosomal rearrangements. The rearrangements we identified were: Firstly, parts of chromosomes NC_036450.1, NC_036459.1, and NC_036466.1 in platyfish made up a single chromosome, Ne4 in Atlantic cod (Figure S5). Secondly, parts of chromosomes NC_036450.1, and 036459.1 in platyfish made up a single chromosome, Ne19 in Atlantic cod (Figure S5). Moreover, between Atlantic cod and John Dory, Ne4 in Atlantic cod is made up of parts of three chromosomes in John Dory OY482846.1, OY482861.1, and OY482865.1 (Figure S6). Furthermore, Ne19 in Atlantic cod is made up of parts of OY482861.1 and OY482865.1 in John Dory. Additionally, Ne12 and Ne22 in Atlantic cod, made up one chromosome in John Dory. Lastly, John Dory OY482846.1, is made up of parts of Atlantic cod Ne4 and Ne21 (Figure S6). Taken together, for most of the chromosomes, chromosomal architecture seems to be fairly conserved in Atlantic cod compared to outgroup species. However, for the rearrangements that are identified, both fissions, fusions, and translocations must have occurred to result in the current chromosomal organization observed in Atlantic cod.

In comparisons carried out between the fish species of the present study, chromosomes Ne4 and Ne19 in Atlantic cod corresponded to one homologous chromosome each in all comparisons, except in European hake where Ne4 in Atlantic cod was found to correspond to parts of Mm1, Mm2, and Mm15 (Figure S7A), and in burbot where Atlantic cod Ne4 was found to be homologous to half of Ll23 and a large part of Ll1 (Figures S7B). Our results suggest that the fusion of chromosomes leading to Ne19 in Atlantic cod (and homologous chromosomes in the other species) either took place before the split of Merlucciidae — since there is a 1:1 relationship between the homologous chromosomes of the other species investigated presently, then a subsequent fission event took place in the lineage of Merlucciidae, or a fusion event occurred after the splitting of Merlucciidae, in the lineage of Gadidae and Lotidae. For Atlantic cod Ne4 our results indicate that a fusion event has occurred after the splitting of Lotidae and Merlucciidae, in the lineage of Gadidae. Moreover, syntenic relationships of Mm1 and Mm15 (European hake) compared to the corresponding chromosomes in the other codfishes (See Figure 2 and Figure S7A for comparisons to burbot and Atlantic cod, respectively) suggest that complex chromosomal reshuffling characterized by both fissions, fusions and/or translocations likely have taken place specifically in the lineage of European hake, which diverged from the Gadidae ~70 million years ago (see Malmstrøm et al.[5]; Figure 1).

Moreover, in the full characterization of the chromosomal architecture among the six Gadiform fish species, Arctic and polar cod were found to harbor eight vs. five larger chromosomes, respectively, likely a result of ancestral chromosomal fusions (see Results and Discussion main document for more details). Additionally, we detect that some of the other gadids: the Atlantic haddock, burbot, and European hake also contain one or several large chromosomes, all exceeding 40 Mbp in size (see Figure 2). These large chromosomes include Ma1 in Atlantic haddock; Ll12 in burbot; and Mm1, Mm2, and Mm3 in European hake (Figure 2, Figure S7A, and Figure S7B). Intriguingly, in comparisons between Atlantic haddock and polar cod, we found that Atlantic haddock Ma1 and polar cod Bs3 — which we identified as being a fused chromosome in polar cod — were 1:1 syntenic (Figure 2 and Figure S7C). Comparisons between Atlantic haddock and Atlantic cod (as well as burbot) show that Ma1 may have originated by a chromosomal insertion event, of which close to the entirety of one ancestral chromosome homologous to Ne15 and Ll19 has been inserted into a chromosome homologous to Ne3 and Ll16 (Figure 2, Figure S7C and S7D). While for Bs3 in polar cod, an end-to-end fusion of chromosomes homologous to Ne3 and Ne15 seems most likely, with subsequent translocation of a smaller part of the end region of Ne15 to the beginning of Bs3. However, for this fused chromosome (Bs3) an insertion of the entire chromosome homologous to Ne3 into Ne15 cannot be ruled out completely (see Figure 2, Figure S7C, and S7D and Results in the main document for more details). From this, we can conclude that even if Ma1 and Bs3 are seemingly 1:1, these fusion events are likely of independent origins in polar cod and Atlantic haddock.

#### Supplementary Materials and Methods

##### Genome sequencing and library preparation

###### *Polar cod, Arctic cod, burbot, and Atlantic cod (NCC)*

DNA was extracted from a variety of tissue samples depending on what was available for each species (Table S5). To obtain high molecular DNA for PacBio as well as Illumina paired end sequencing libraries, DNA isolation was performed using a high-salt extraction protocol (SOP can be found at https://www.mn.uio.no/cees/english/people/researcher-postdoc/jentoft/sop-038-high-salt-dna-extraction.pdf). PacBio sequencing libraries were prepared using the PacBio 20 kb library preparation protocol (PacBio). Fragmentation of DNA was performed using Megaruptor (Diagenode) and size selection done using BluePippin (Sage Science) with a cut-off length of 10 kb. Libraries were sequenced on a Pacific Biosciences Sequel II instrument (PacBio), using Sequel Polymerase v2.0 and Sequencing Chemistry v.2.1, and loading on the instrument was performed by diffusion. The libraries were sequenced across 12, 11, and 10 SMRT cells, for polar cod, Arctic cod, and burbot respectively with a movie time of 600 minutes.

The Atlantic cod (NCC) genome was sequenced across three rounds and on three PacBio, where the two first libraries were sequenced on Pacific Biosciences RS II instrument using P6-C4 chemistry. A total of 28 SMRT cells were used for sequencing, 15 SMRT cells were sequenced with 240 min movie time, and 5 SMRT cells were sequenced using 360 min movie time. Additionally, eight additional SMRT cells (with a movie time of 600 minutes) were sequenced using the Sequel instrument (PacBio), using Sequel Polymerase v2.0, and Sequencing Chemistry v.2.0 (Table S2).

For polar cod, Arctic cod, and burbot preparation of Illumina 150 bp paired-end (PE) libraries were done using KAPA HyperPrep Plus (Roche), with 4 cycles of PCR. Sequencing was performed across 3 lanes, using the Illumina Hiseq4000 instrument (Illumina). For polar cod, Arctic cod, and Atlantic cod, high molecular DNA for 10X library preparation was isolated from gill or spleen (Table S5) using the Nanobind CBB BIG DBA kit (Circulomics Inc.). 10X library preparation was performed following the Chromium Genome Reagent kit V2 (10X Genomics) user guide using the 10X listed kits as well as suggested QC kits and methods including Chromium™ Genome Chip Kit v2, 48 rxns, Chromium™ Genome Library Kit & Gel Bead Kit v2, 16 rxns, Chromium™ Genome Library Kit V2, 16 rxns -120255, Chromium™ Genome Gel Bead, 16 rxns -120214, and Chromium™ i7 Multiplex Kit, 96 rxns. The 10X libraries were sequenced as 150 bp paired end reads on the Illumina HiseqX Instrument (Illumina). For polar cod, Arctic cod and Atlantic cod (NCC), gill filaments and spleen were used as input tissue (Table S5) with the Arima Hi-C Kit (Arima Genomics) to obtain crosslinked and proximally ligated DNA-PLD. For burbot Omni-C (Dovetail Genomics) was utilized for crosslinking DNA. 150 bp paired end Hi-C library preparation was performed using KAPA Hyper Prep Kit (Roche) creating Illumina library from the PLD, and sequenced on the Illumina HiseqX Instrument (Illumina).

***European hake***

Isolation of DNA from European hake was done from blood, using the nucleated blood protocol by the Circulomics Nanobind BIG DNA kit (Circulomics Inc.). The PacBio libraries were prepared using the Pacific Biosciences protocol for HiFi library prep using SMRTbell® Express Template Prep Kit 2.0. Two libraries were prepared, and 7.5 µg DNA was fragmented into 15-20 kb fragments using Megaruptor 3. A total of 5 µg of fragmented DNA was used for library prep. Final library was size selected using BluePippin with a 10 kb cut-off. Library was sequenced on two 8M SMRT cells on a Sequel II instrument using Sequel II Binding kit 2.2 and Sequencing chemistry v2.0. Loading was performed by adaptive loading, with a movie time of 30 hours. Hi-C preparation was done using the Dovetail Omni-C kit provided by Dovetail genomics. Library preparation followed the "Omni-C proximity Ligation Assay, version 1.0". The final library was sequenced on a S4 flow cell utilizing the 2*150 bp PE mode on an Illumina Novaseq sequencer. DNA isolation, HiFi sequencing, and Hi-C sequencing were performed by the Norwegian Sequencing Centre (NSC).

***Atlantic haddock***

Isolation of DNA from haddock was done from blood (Table S5), using the Nanobind HMW Tissue DNA kit (Circulomics Inc.). Library was prepared using Pacific Biosciences Express library preparation protocol without any fragmentation of the sample prior library prep. Size selection of the final library was performed using BluePippin using 15 kb cut-off. The library was sequenced on 6 SMRT cells using Sequel II Binding kit 2.0 and Sequencing chemistry v2.0 with 15 hours movie time. The Hi-C library was prepared following instructions in the Arima library user guide.

Around 4.4μg proximally ligated DNA was sheared using Covaris tubes on a Covaris E220 instrument. Following size selection, 383ng was used in the biotin enrichment step. Illumina unique indexing adaptors were used for the ligation. Library was amplified with 6 cycles of PCR, purified, and checked on a fragment analyzer (FA) using the NGS kit. Library concentration was controlled with qPCR (KAPA Quantification kit). An additional Illumina library was prepared based on the isolated DNA from the Circulomics DNA protocol, with 1000ng gDNA as input in the Kapa Hyper prep PCR-free workflow. The same quality checks as for Hi-C were performed here.

***De novo* genome assembly for six Gadiform fishes**

To construct the primary assembly for Arctic cod, Atlantic cod (NCC), and burbot, PacBio Sequel long reads were assembled using Flye v2.4[6]. The option "minimum overlap" was set to 2.5 kbp for all three primary assemblies. The presence of duplicated contigs was evaluated using purge dups v.1.0.1[7], and removal of duplicated contigs in the primary assembly of Arctic cod, as well as burbot, was performed using purge dups. In order to scaffold the primary contigs into chromosome length scaffolds, we leveraged a combination of 10X linked and Arima Hi-C linked Illumina PE sequencing data for Arctic cod and Atlantic cod (NCC).

To scaffold the primary assembly of Arctic cod and Atlantic cod (NCC), Illumina reads from a 10X sequencing library were aligned to the primary assemblies scaffolded/linked using Scaff10X v4.2[8]. Subsequently, Illumina Hi-C sequence data were mapped to the 10X scaffolded assembly using Juicer v1.5.6[9]. For the burbot, Hi-C OmniC Illumina PE reads were mapped directly onto the Flye primary assembly using bwa mem v0.7.17[10] and scaffolded by Juicer v1.6[9].

Contigs could then be grouped, arranged, and anchored to near chromosomal length scaffolds using 3D-DNA v180922[11]. The 3D-DNA draft assemblies and Hi-C contact maps for each species produced by Juicer were visualized and manually inspected in the Juicebox program-suite v1.11.08[12]. During manual curation of Hi-C contact maps, we discovered two obviously erroneous linked super-scaffolds within the Atlantic cod (NCC) as well as the burbot assembly, which had low Hi-C linkage between them, and which likely make up two separate chromosomes. These were manually split into two super-scaffolds, and we used the 3D-DNA post-review script to correct the assemblies (https://github.com/aidenlab/3d-dna/blob/master/run-asm-pipeline-post-review.sh). After scaffolding the genome assemblies for all three species, we performed additional rounds of polishing to close any remaining gaps and improve the base level accuracy of the assemblies. For long-read polishing, we utilized

PacBio sequencing data and aligned the reads to the Hi-C scaffolds using pbmm2 v1.2.1[13]. polishing was performed using gcpp v1.9.0[13]. Lastly, two rounds of short-read polishing were performed by aligning Illumina paired-end reads with Minimap v2.155[14] and subsequent base calling, done using Freebayes v1.3.2[15]. Finalized genome assemblies were evaluated using metrics calculated by the Assemblathon_stats[16] script, completeness of each assembly was assessed using the Benchmarking Universal Single-Copy Orthologs (BUSCO) software v5.0.0[17] as well as by aligning and visualizing the final assemblies against the chromosome level assembly of Atlantic cod (gadMor3.0) [4] using the D-GENIES homepage[18].

The European hake genome was assembled using a combination of PacBio HiFi long reads and Hi-C data. First, HiFi reads were assembled into a primary assembly using Flye v2.9[6] with default settings, and the parameter genome size was set to 650 Mb. HiFi reads were then mapped against the assembly using Minimap2 v2.17[14] to generate a read-depth histogram to assess duplication levels. Purge Haplotigs v1.1.2[19] was used to identify and purge any potentially duplicated contigs. Reads were flagged and purged using the read-depth information from the mapping step and cutoffs were manually selected based on the read-depth histogram.
Following, the purged Flye draft assembly was scaffolded using Juicer v1.22.01[9] and the 3D-DNA pipeline[11] by first aligning Hi-C data to the purged primary assembly using the Juicer pipeline. Next, the alignment file was run through the 3D-DNA pipeline to produce a candidate near chromosome-length draft genome assembly. Thereafter, the draft genome assembly was visualized using Juicebox v1.11.08[12] and manually curated for any misassembles. Manual curation included splitting one super-scaffold into two, due to weak Hi-C contact points between scaffolds. The manually curated assembly was processed through the 3D-DNA pipeline one last time, incorporating the changes made within Juicebox.

Atlantic haddock was assembled using a combination of PacBio long reads and chromosome confirmation Hi-C reads. The PacBio reads were first assembled using Flye v2.9 with default settings. The PacBio reads were mapped to the Flye draft assembly with minimap v2.22 and then purge_dups v1.2.5 was applied. Next, Hi-C reads were mapped against the purged genome assembly with bwa mem v0.7.17 with the options -5SPM. SAMtools v1.11[20] was used to deduplicate the mapped reads. YaHS v1.1a[21] was used to scaffold the purged genome assembly. Next, pbmm2 v1.9 was used to map the PacBio reads to the scaffolded assembly, and gcpp v1.0.0 was used to polish the assembly. Further, two rounds of polish using Illumina reads mapped with bwa mem v0.7.17 and processed with Freebayes v1.3.6. FCS-Adaptor v0.2.2 (https://github.com/ncbi/fcs) was run on the polished assembly, and any adaptor sequences found were masked using BEDtools v2.30.0 maskfasta[22]. FCS-GX v0.2.2 (https://github.com/ncbi/fcs) was used to search for contamination. If a contaminant was found at the start or end of a sequence, the sequence was trimmed using a combination of SAMtools faidx[20], BEDtools complement and BEDtools getfasta. If the contaminant was internal, it was masked using BEDtools maskfasta. The assembly was manually curated using the GRIT rapid curation suite[23] and the PretextView v.0.2.5 (https://github.com/wtsi-hpag/PretextView, last accessed September 27, 2023). Genome assembly metrics were calculated using the Assemblathon_stats[16] script, completeness of each assembly was assessed using the Benchmarking Universal Single-Copy Orthologs (BUSCO) software v5.0.0[17] as well as by aligning and visualizing the final assemblies against the chromosome level assembly of Atlantic cod (gadMor3.0)[4] using the D-GENIES homepage[18].

**RNA sequencing for annotation**

Total RNA was extracted from spleen, liver and gonad for polar cod and Arctic cod. All tissues were stored at -80 °C from collection till arrival and processing at University of Oslo, Norway. RNA isolation was performed using the RNeasy mini kit (Qiagen) following the manufacturer's instruction. RNA isolate was normalized, and 2µg was used as input for the Illumina TruSeq mRNA stranded kit (Illumina) and prepped on a Perkin Elmer Sciclone NGSx liquid handler system (Perkin Elmer). Samples were indexed using Uniqued dual indexing. Final libraries were QCed on a fragment analyzer system using the standard sensitivity NGS kit from AATI (Agilent) and quantified using the KAPA Library quantification kit for Illumina (Roche). The library pool was sequenced on an Illumina Hiseq4000 instrument using 2*150bp PE read mode (Illumina).

**Genome annotation**

***Norwegian coastal cod (NCC), Arctic cod, and polar cod***

A library of repeated elements was created as described in Tørresen et al. [24]. Briefly, RepeatModeler v1.0.8[25], LTRharvest[26], part of genometools v1.5.7[27], and TransposonPSI[28] were used in combination to create a set of putative repeats. Elements with a match only against an UniProtKB/SwissProt database[29] and not against the database of known repeated elements included in RepeatMasker were removed. The remaining elements were classified and combined with known repeat elements from RepBase v20150807[30].

HISAT v2.1.0[31] was used to map all RNA-seq data to the assembly. For NCC we downloaded the following datasets from SRA: PRJEB12487, PRJEB31396, PRJNA256972, PRJNA277848, and PRJNA277848. Portcullis v1.2.0[32] was run on the mapped RNA-seq reads to generate a catalog of good junctions. These junctions were used in a second round of HISAT2. Mikado v2.0rc6[33] created a set of the best transcripts from the second round of HISAT2. Predicted proteins from zebrafish (*Danio rerio*), Atlantic cod (gadMor1 and gadMor3), channel bull blenny (*Cottoperca gobio*), herring (*Clupea harengus*), Northern pike (*Esox lucius*), denticle herring (*Denticeps clupeoides*) and *Triplophysa tibetana* was downloaded, and used in ProtHint v2.2.0[34] to generate hints for BRAKER. BRAKER v2.1.5[35] was run with the hints from ProtHint and mapped reads from the second round of HISAT2. GenomeThreader v1.7.1[36] was used to map UniProtKB/Swiss-Prot Release 2019_10[29] of 13-Nov-2019 proteins to the genome assembly. The funannotate v1.7.0 [37] mask command was used to mask the genome assembly with the repeat library, before funannotate train was run with all RNA-seq data to train AUGUSTUS v3.3.3[38] The training results from funannotate train, proteins from the fishes and UniProtKB/Swiss-Prot, GeneMark gene models from BRAKER, and gene models from Mikado and GenomeThreader were used as input to funannotate predict. InterProScan v5.34-73.0[39] was run on the predicted proteins from funannotate predict, and funannotate annotate was used to integrate those results into a final predicted set of genes.

***Atlantic haddock, burbot, and European hake***

AGAT v1.0[40] agat_sp_keep_longest_isoform.pl and agat_sp_extract_sequences.pl were used on the zebrafish assembly and annotation to generate one protein (the longest isoform) per gene. Miniprot v0.5[41] was used to align the proteins to the curated assemblies. UniProtKB/Swiss-Prot release 2022_03[29] in addition to the vertebrata part of OrthoDB v10[42] were also aligned separately to the assemblies. RED v2018.09.10 was run via redmask (https://github.com/nextgenusfs/redmask) on the assemblies to mask repetitive areas[43]. GALBA ((https://github.com/Gaius-Augustus/GALBA, commit: f4aaeca) was run with the zebrafish proteins using the miniprot mode on the masked assemblies [41,44–47]. The funannotate-runEVM.py script from Funannotate v1.8.13[48] was used to run EvidenceModeler v1.1.1[49] on the alignments of zebrafish proteins, UniProtKB/Swiss-Prot proteins, vertebrata proteins, and the predicted genes from GALBA. The resulting predicted proteins were compared to the protein repeats that funannotate distributes using DIAMOND v2.0.15[45] blastp and the predicted genes were filtered based on this comparison using AGAT. The filtered proteins were compared to the UniProtKB/Swiss-Prot release 2022_03 using DIAMOND blastp to find gene names and InterProScan v5.47-82[39] was used to discover functional domains. AGATs[40] agat_sp_manage_functional_annotation.pl was used to attach the gene names and functional annotations to the predicted genes.

##### *Long-read assembly of mitochondrial genomes*

Mitochondrial genomes (mitogenomes) were generated using either of two methods depending on the input data available. Species sequenced using PacBio continuous long reads (CLR) and which had Illumina PE short reads available were assembled using the mitoVGP v2.0 pipeline[1]. The MitoHiFi v2.2[2] pipeline was applied to assemble the mitogenomes of the Atlantic cod (NEAC) straight from HiFi data. MitoHiFi can either assemble mitogenomes from raw HiFi reads or search already assembled contigs to extract potential mitogenomes. In addition to the assembly from raw reads, the European hake draft purged Flye assembly was used to search for and retrieve any potentially assembled mitogenomes. Raw reads that map to the mitogenomes are extracted as part of the assembly pipeline of MitoHiFi and MitoVGP. These reads were mapped back to the assembled mitogenomes using pbmm2 v1.9.0[13], a Minimap2 SMRT wrapper for PacBio data, to assess read depth distribution, and read depth was measured per base using SAMtools v1.14[20]. The mitogenomes were afterward annotated and visualized using the online tool MitoFish v3.74[50,51]. Lastly, after assembly and annotation mitogenomes for all seven species were manually inspected using the Integrative Genomics Viewer (IGV) v2.9.4[52].

#### Phylogenomic placement of Arctic cod

##### *Mitochondrial phylogeny*

For the codfishes with sequenced genomes in this study, except for Atlantic haddock which was accessed from NCBI accession: NC_007396.1, mitochondrial protein-coding genes (PCGs) identified by MitoFish were aligned using MAFFT v7.453[53] and manually inspected and corrected for reading frame shifts before they were concatenated with PhyKIT v1.11.7 create_concat[54] to produce a supermatrix. A maximum likelihood (ML) tree was inferred using IQ-Tree2 v2.2.0[55] and ModelFinder Plus (MFP)[56] was used to search for the best substitution model under the Bayesian information criterion (BIC). Branch support was calculated using 1,000 replicates of ultrafast bootstrap approximation (UFBoot)[57].

A Bayesian approach to tree inference was done in BEAST v2.6.7[58] using the same concatenated supermatrix. BEAST Model Test (bmodeltest)[59] was used to infer the substitution model from the namedExtended list of substitution models. A strict clock was applied using the birth-death model as prior and the analysis was run with a chain length of 100,000,000, with sampling done every 1,000 iterations for a total of 100,000 trees. Convergence was assessed using Tracer v1.7.2[60] and TreeAnnotator was used to produce the consensus tree with a burn-in of 10% and the target tree set to maximum clade credibility.

Additionally, phylogenetic analysis was conducted using the complete mitogenomes to compare the effects of including all data vs only the PCGs on topology inference. MitoFish outputs reordered mitogenomes, i.e., all start at the same position (tRNA-phe). Therefore, MitoFish annotated mitogenomes were aligned using MAFFT v7.453[53], and gaps were trimmed using ClipKIT v1.3.0[61] with the smart-gap approach. The smart-gap strategy was selected because it should work better with alignments that contain taxa that span both shallow and deep evolutionary timescales, appropriate for the taxa included in this study (ClipKIT documentation:https://jlsteenwyk.com/ClipKIT/performance_assessment/index.html #smart-gap). Tree inference for the complete mitogenomes followed the same steps as the PCGs for both ML and Bayesian analysis.

### Supplementary Figures

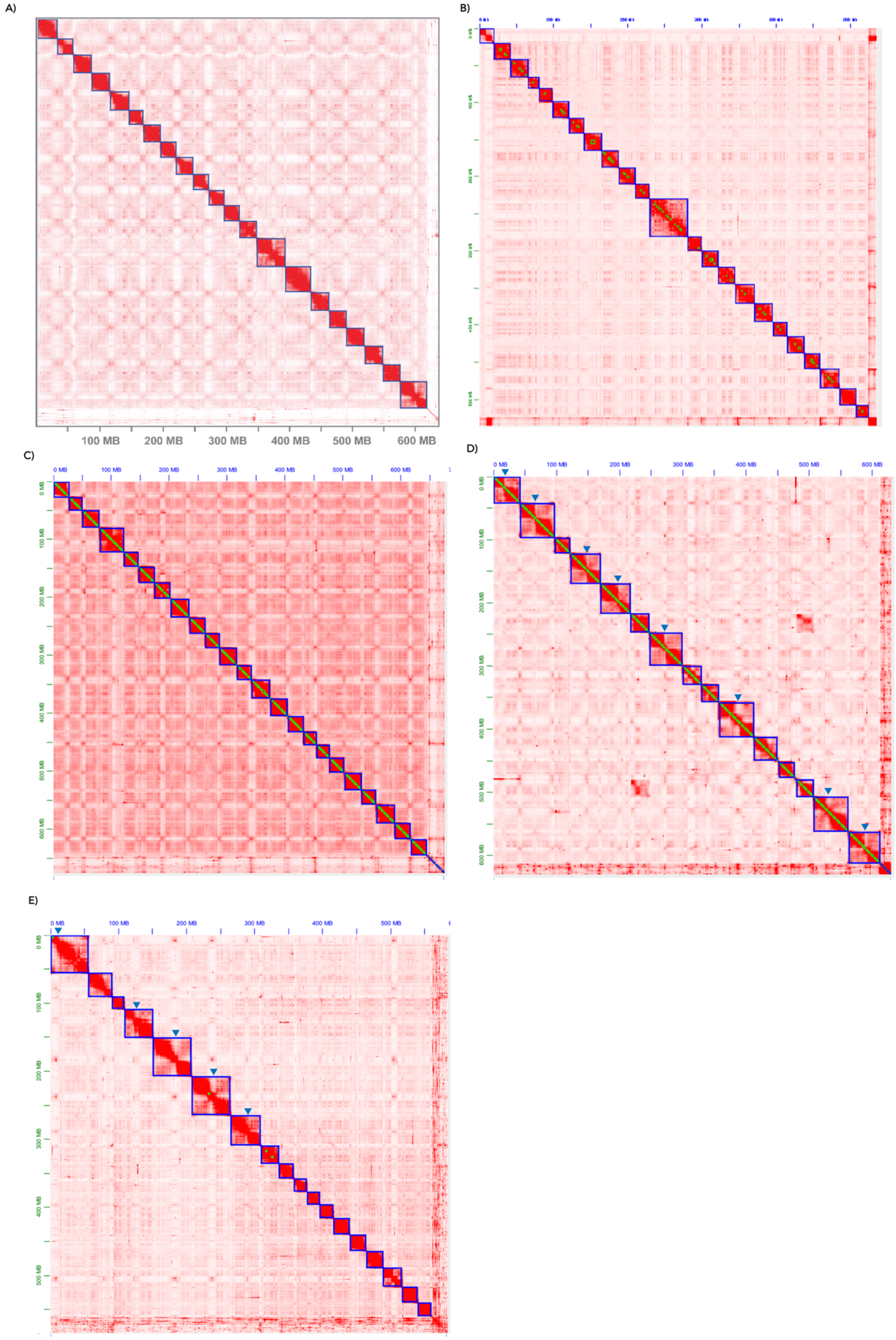

F)

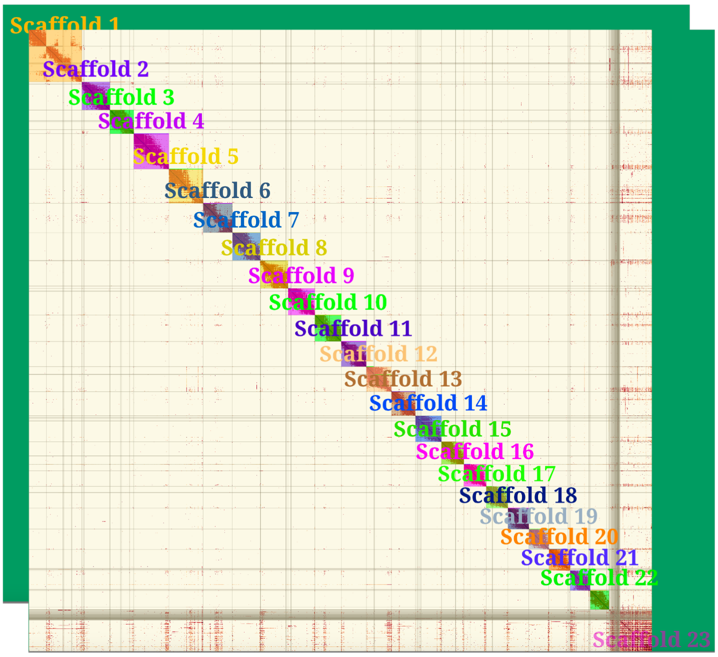

**Figure S1.** Hi-C contact points are shown as heatmap on super-scaffolds of each final genome assembly of the six Gadiform fishes. Regions assumed to make up centromeres annotated with blue triangles for the fused chromosomes in Arctic cod and polar cod. **A)** European hake **B)** burbot**, C)** Atlantic cod (NCC), **D)** Arctic cod, **E)** polar cod, and **F)** Atlantic haddock.

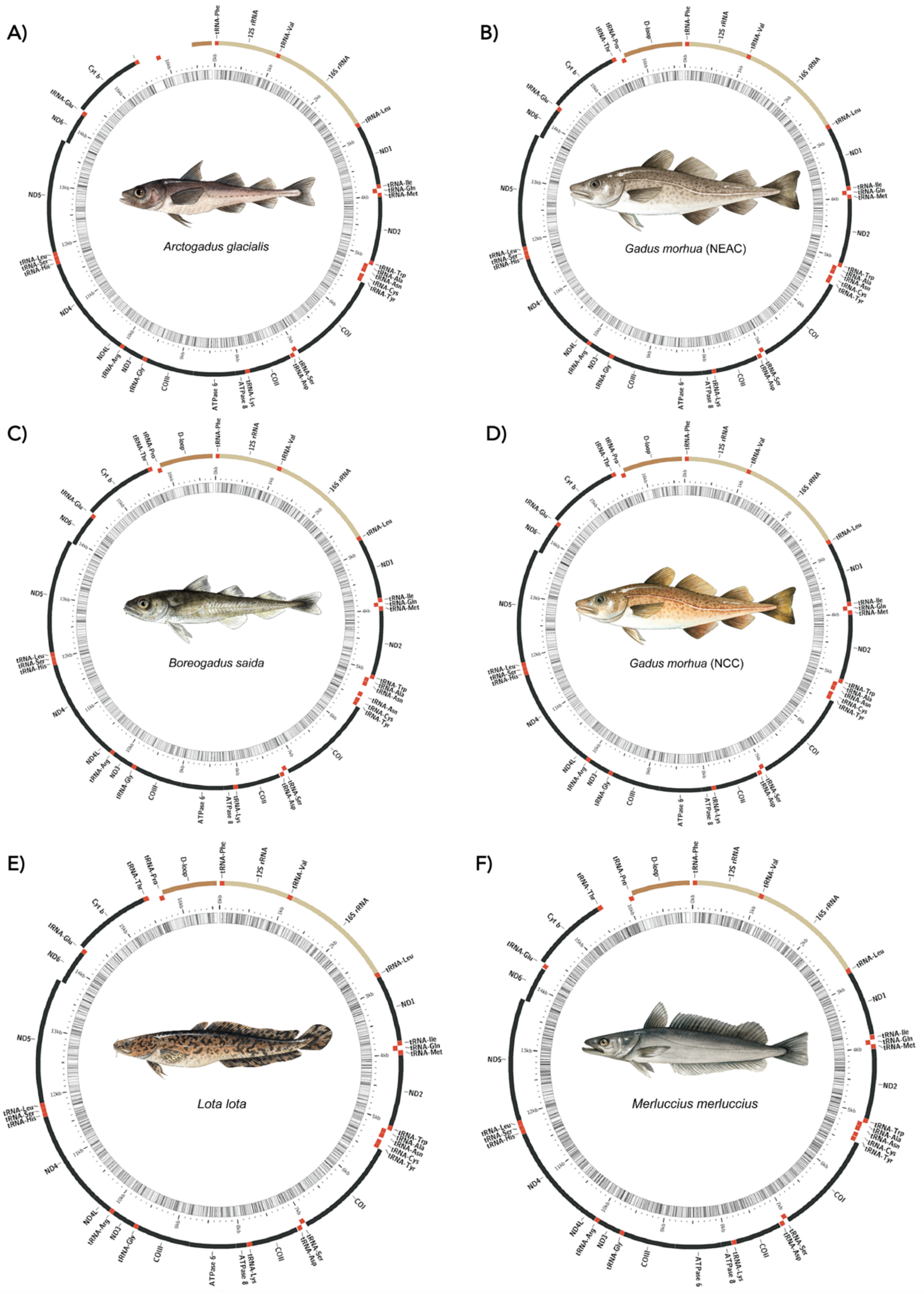
A)
Arctogadus glacialis
B)
Gadus morhua (NEAC)
C)
Boreogadus saida
D)
Gadus morhua (NCC)
E)
Lota lota
F)
Merluccius merluccius

G)

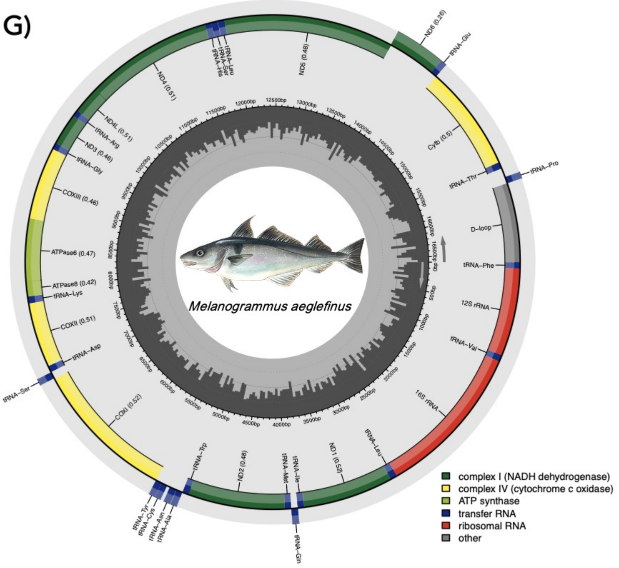

**Figure S2.** Mitofish annotated mitogenomes for **A)** Arctic cod, **B)** Atlantic cod (NEAC), **C)** polar cod**, D)** Atlantic cod (NCC**), E)** Burbot**, F)** European hake and **G)** Atlantic haddock. In **A-F)** the inner band represents GC-content. All fish illustrations by Alexandra Viertler.

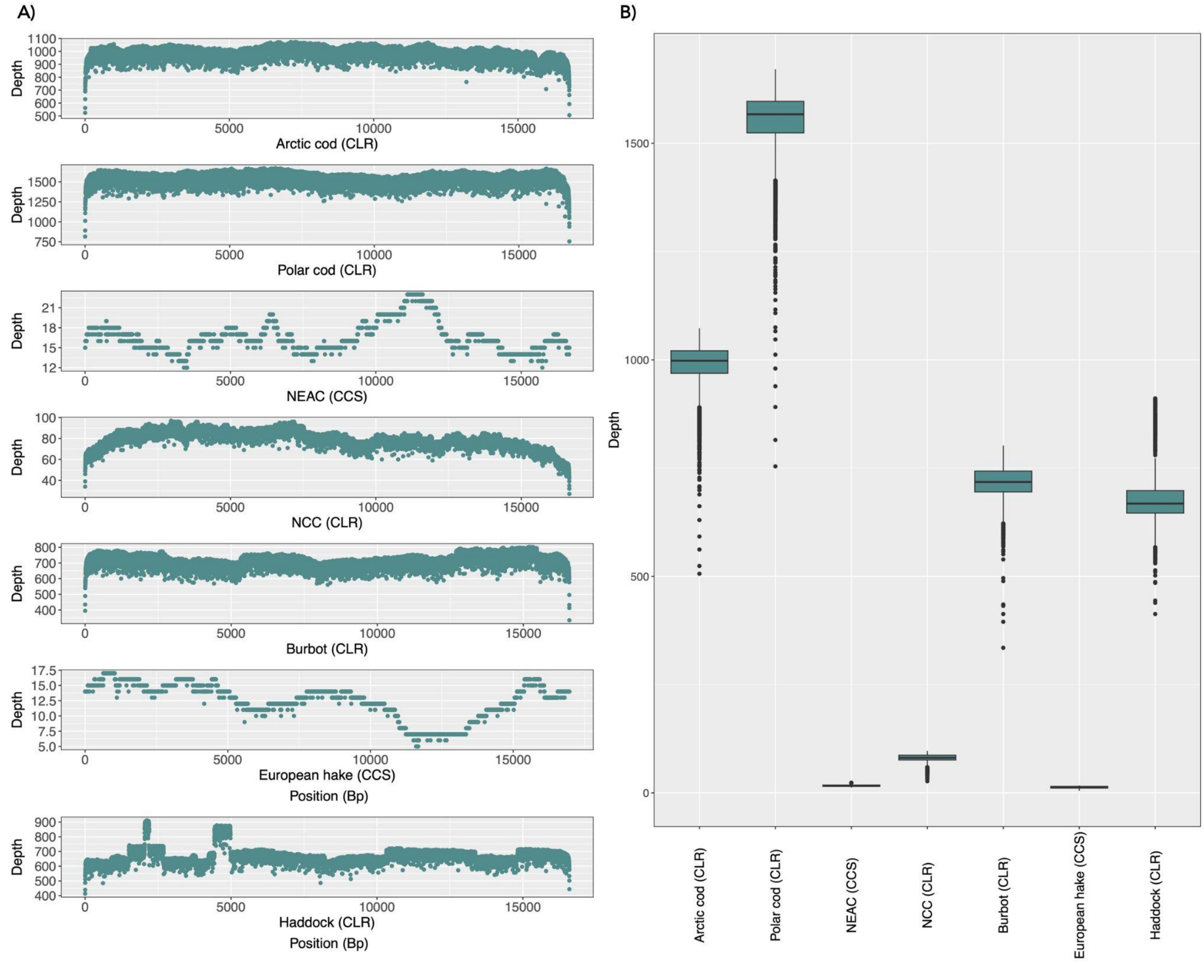

**Figure S3.** Depth distribution of PacBio long-reads mapped against the assembled mitogenomes. A) Depth of the reads along the mitogenomes per base and B) Depth visualized as boxplots per assembly.

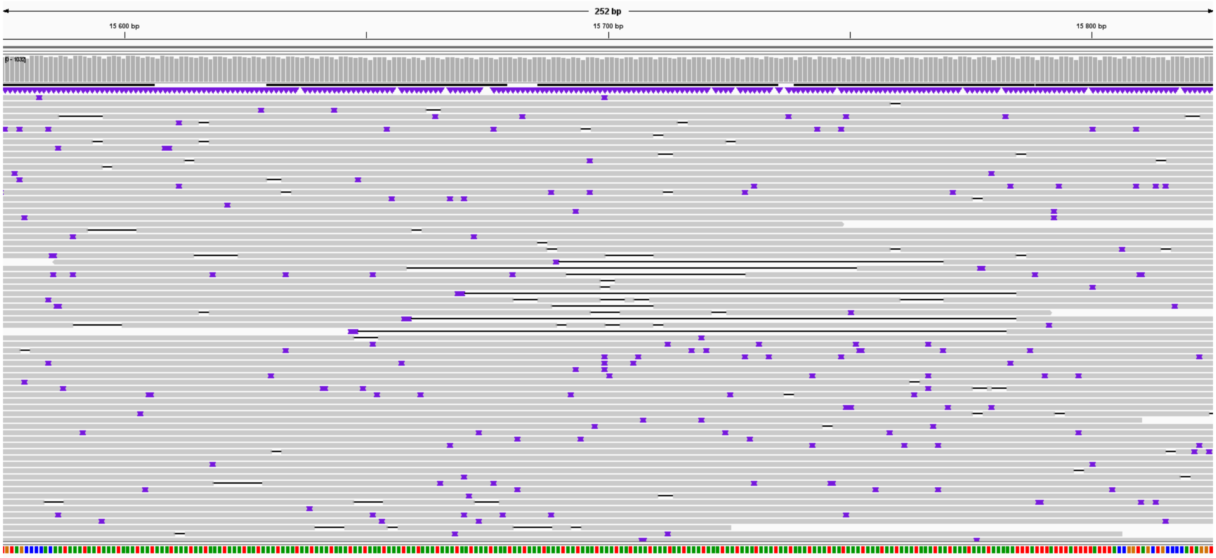

**Figure S4.** Examples of potential heteroplasmic reads along a repeat within the Arctic cod mitochondrial genome from position 15575 - 15825 visualized in

Integrative Genomics Viewer[62]. Purple points and black bars represent insertions or deletions, respectively.

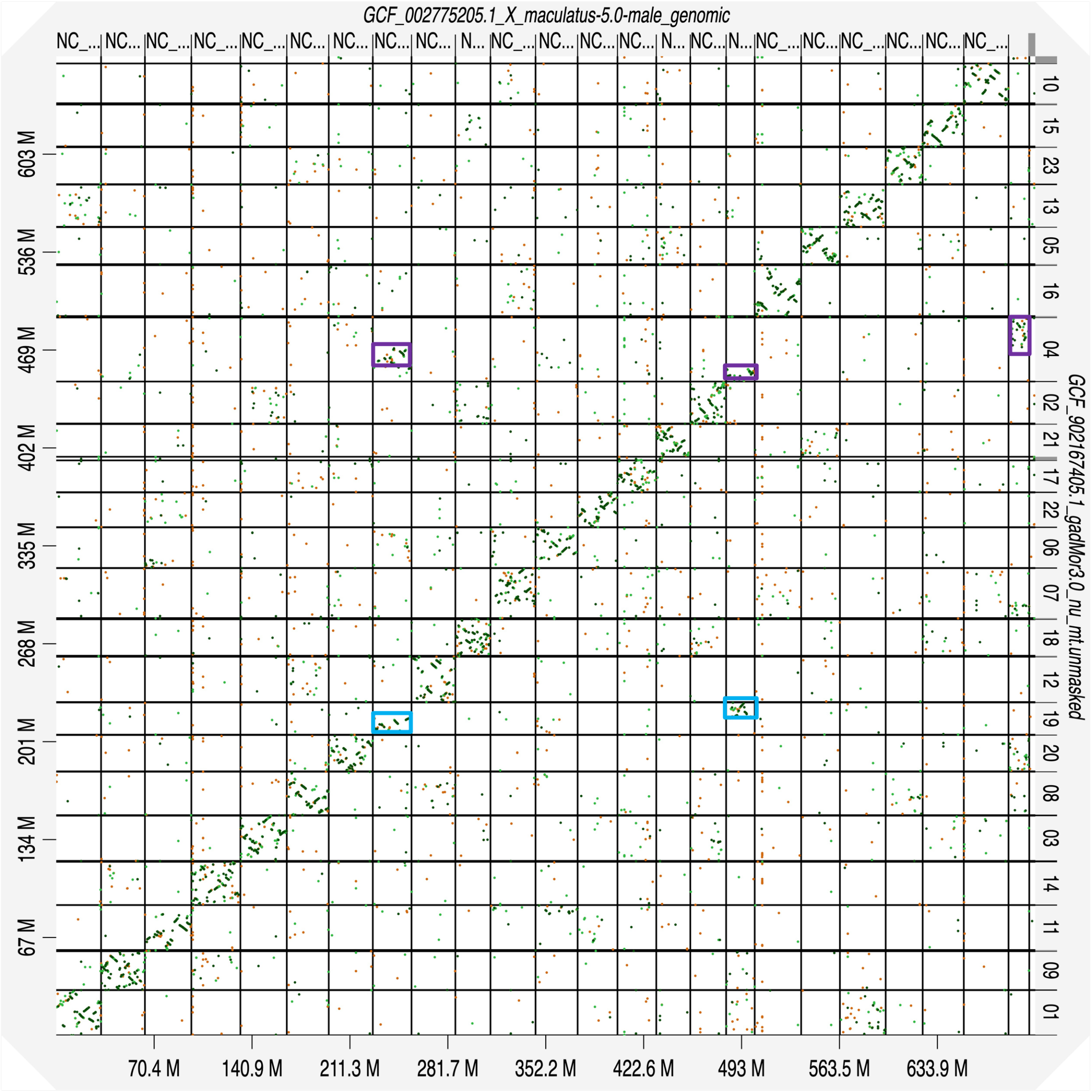

**Figure S5.** Dotplot between Atlantic cod (NEAC) and platyfish (n=24) genome assembly reveals mostly syntenic relationships between chromosomes. Two possible fused chromosomes in Atlantic cod (LG04 and LG19) compared to platyfish marked with squares.

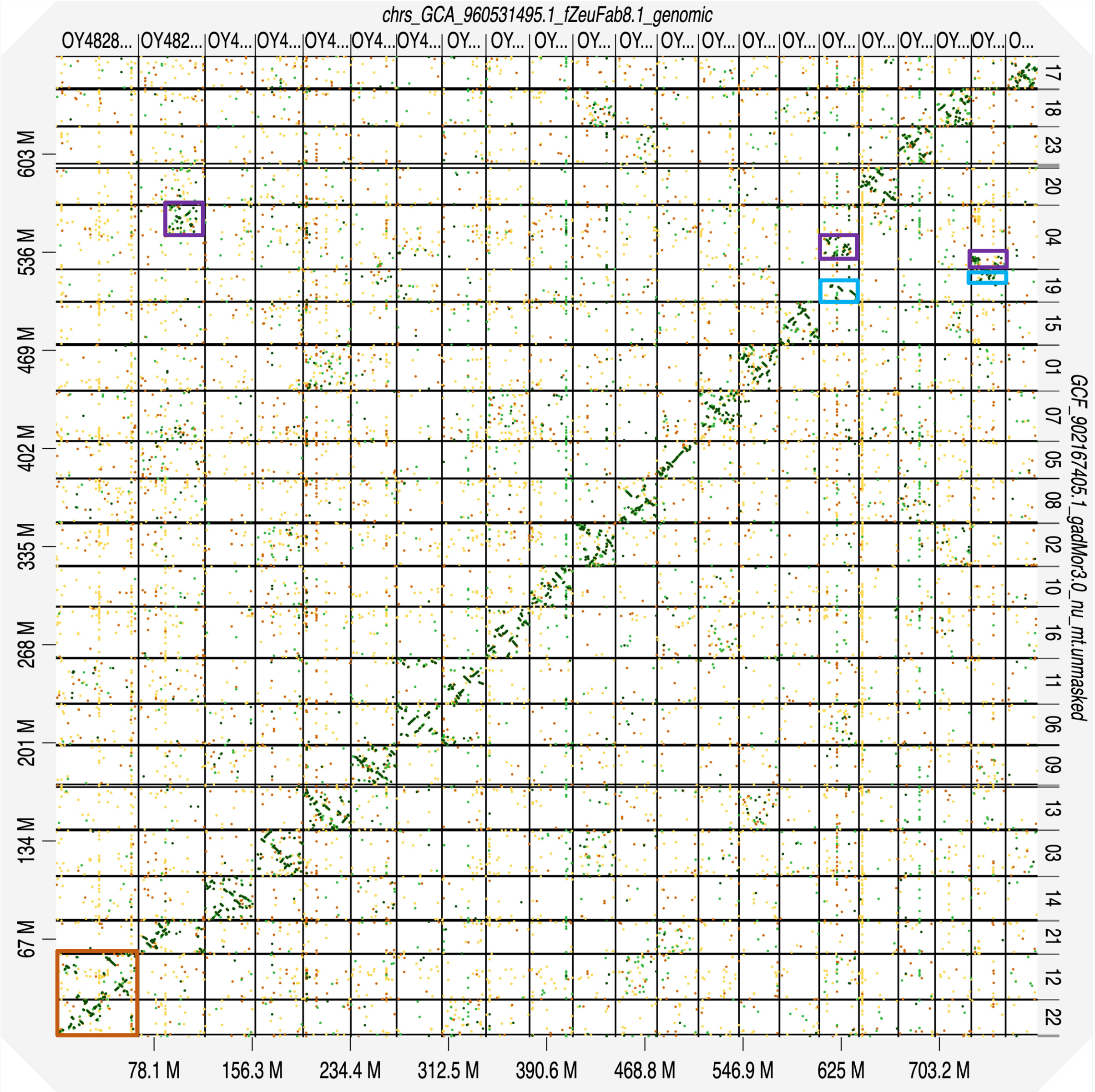

**Figure S6.** Dotplot between Atlantic cod (NEAC) and John Dory (n=22) genome assembly. Two possible fused chromosomes in Atlantic cod (LG04 and LG19) compared to John Dory marked with squares. A fission (Atlantic cod) or fusion (John Dory) is marked in orange.

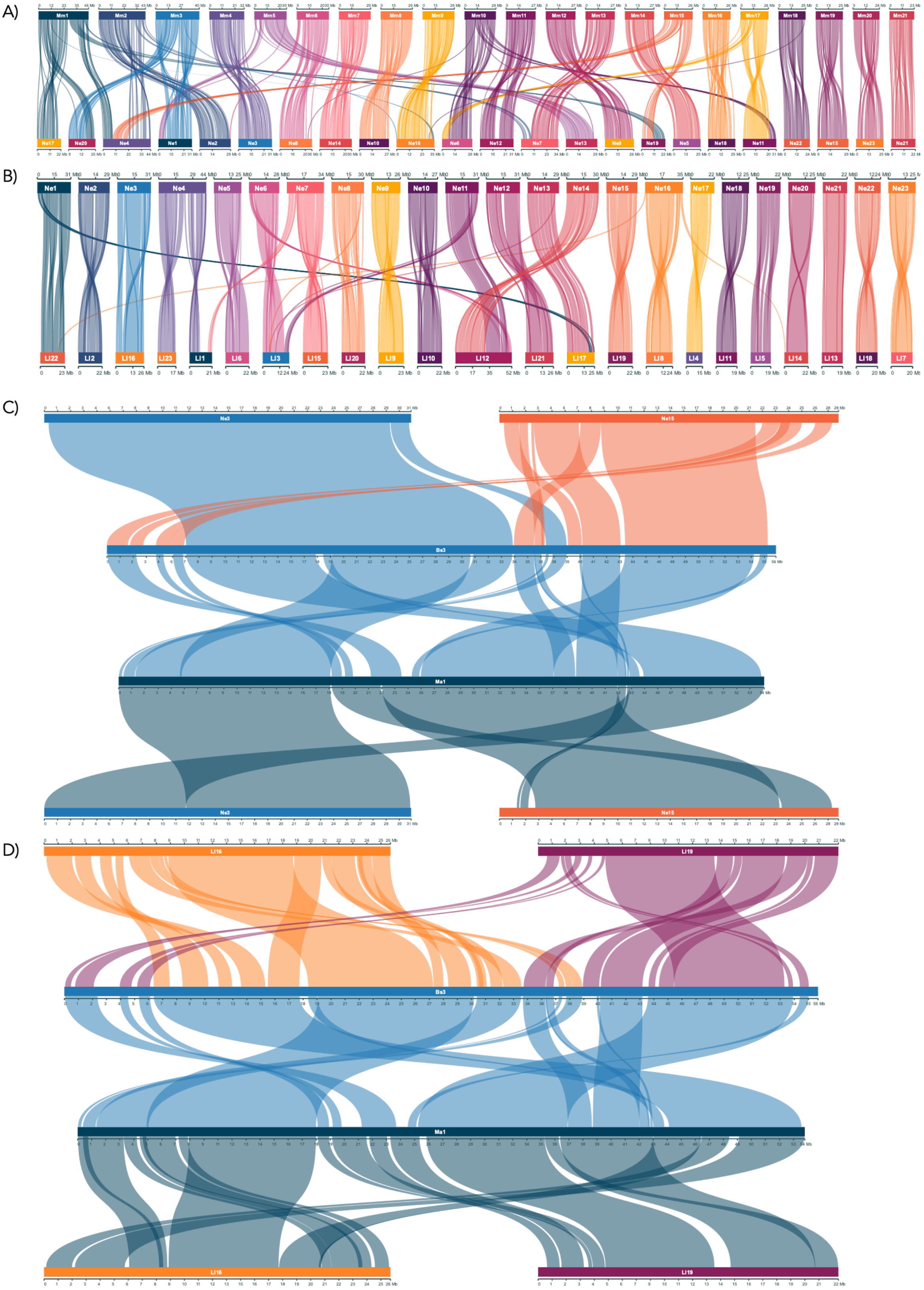

**Figure S7.** Chromosomal synteny based on gene order between Atlantic cod (NEAC) and **A)** European hake, and **B)** burbot. Chromosomal synteny between homologous chromosomes polar cod Bs3 and Atlantic haddock Ma1, as well as homologous chromosomes in **C)** Atlantic cod (Ne3 and Ne16), and **D)** burbot (Ll16 and Ll19).

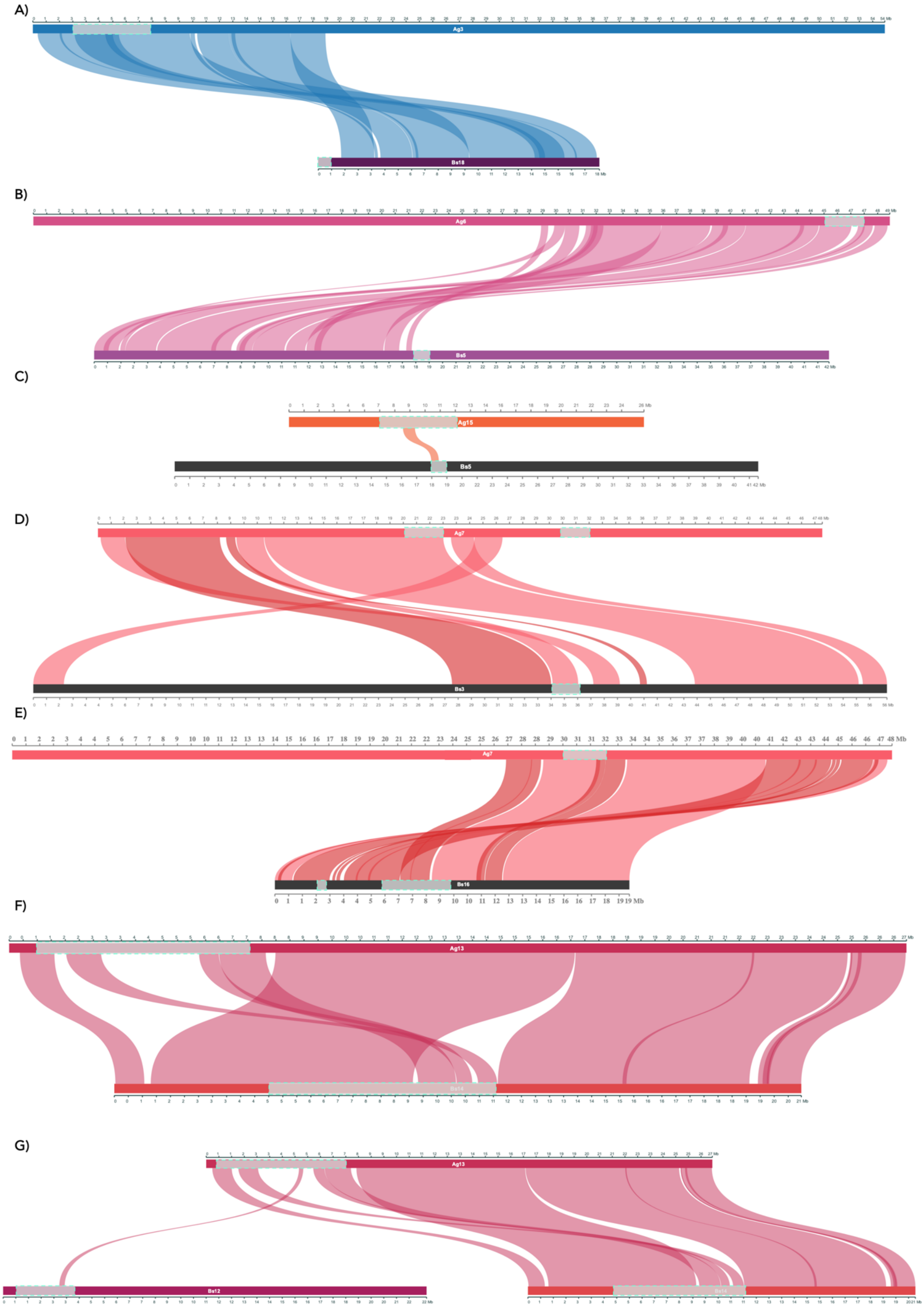

**Figure S8.** Chromosomal synteny between homologous chromosomes harboring species specific inversion in Arctic cod and polar cod. **A)** Ag3 - Bs10 with no overlap detected between inversions. The beginning of Bs18 is homologous to the end of Ag4, harboring no inversion in Arctic cod (Figure 3A). **B)** Ag6 - Bs5, no overlap detected between inversions. **C)** Between the homologous regions harboring inversions on Arctic cod Ag15 and polar cod Bs5 a small overlapping genomic region was detected. **D)** No overlap between inversions was observed between Ag7 - Bs3. **E)** The inversions on Arctic cod Ag7 (~30 - 32 Mb) and polar cod Bs16 (~6 - 9.8 Mb) were found to share one breakpoint region**. F)** We found that the homologous genomic regions of inversions on Arctic cod Ag13 (~1 - 7 Mb) and polar cod Bs14 (~5 - 11 Mb) overlap to some extent. **G)** We additionally found a small genomic region of the inversion on Arctic cod Ag13 to be homologous to the region within a polymorphic inversion on Bs12 in polar cod.

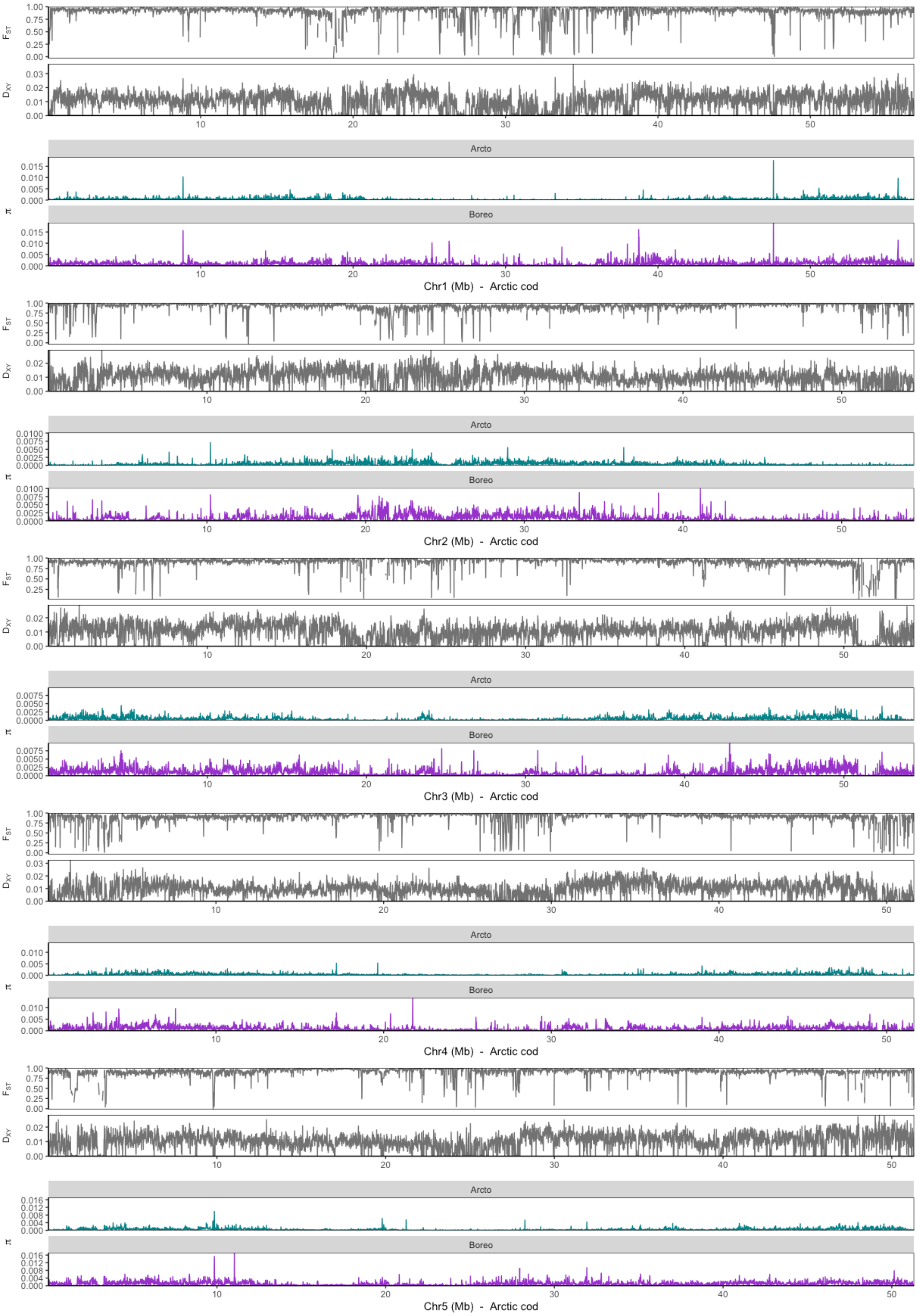
F_ST
D_XY
Arcto
Boreo
π
Chr1 (Mb) - Arctic cod
Chr2 (Mb) - Arctic cod
Chr3 (Mb) - Arctic cod
Chr4 (Mb) - Arctic cod
Chr5 (Mb) - Arctic cod

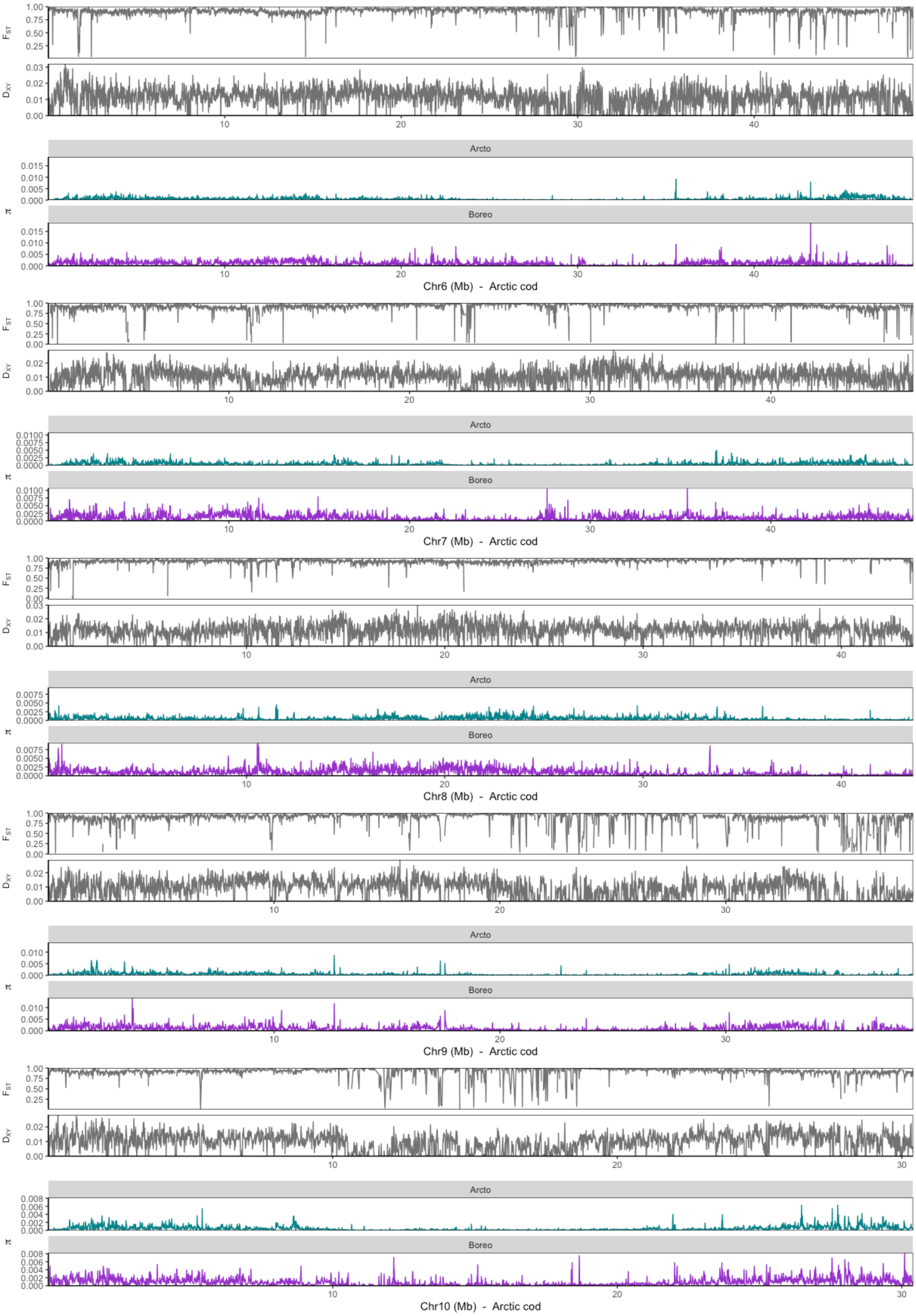
F_ST
D_XY
π
Arcto
Boreo
Chr6 (Mb) - Arctic cod
Chr7 (Mb) - Arctic cod
Chr8 (Mb) - Arctic cod
Chr9 (Mb) - Arctic cod
Chr10 (Mb) - Arctic cod

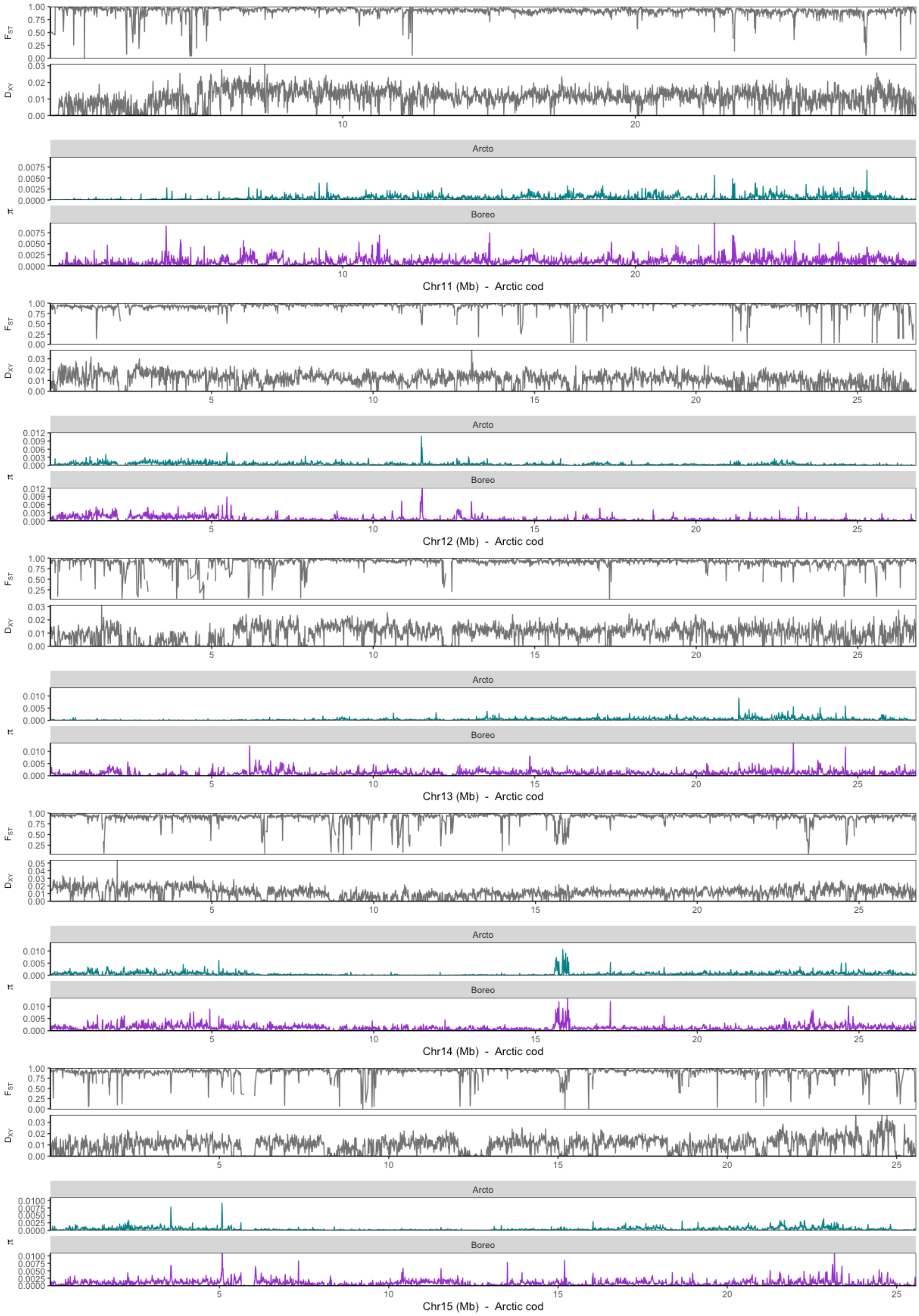

**Figure S9.** Intraspecies genetic differentiation and nucleotide diversity estimated using pixy[63] between Arctic cod and polar cod using the Arctic cod genome assembly as a reference, for chromosomes 1 – 15 (i.e., Ag1-15).

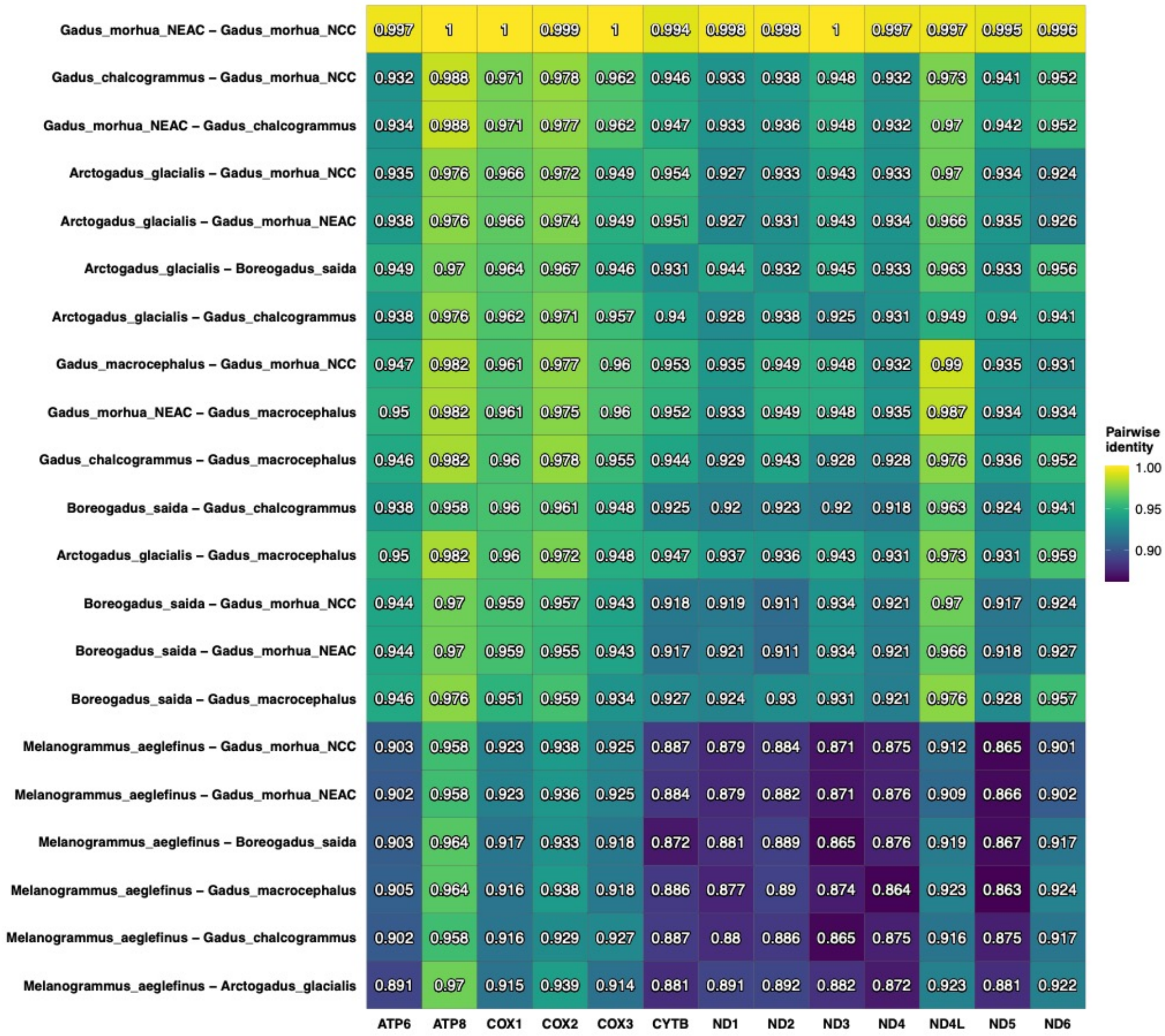

**Figure S10.** Pairwise identity calculated with PhyKIT[54] for each mitochondrial gene between Arctic cod, polar cod, NEAC, NCC and Atlantic haddock.

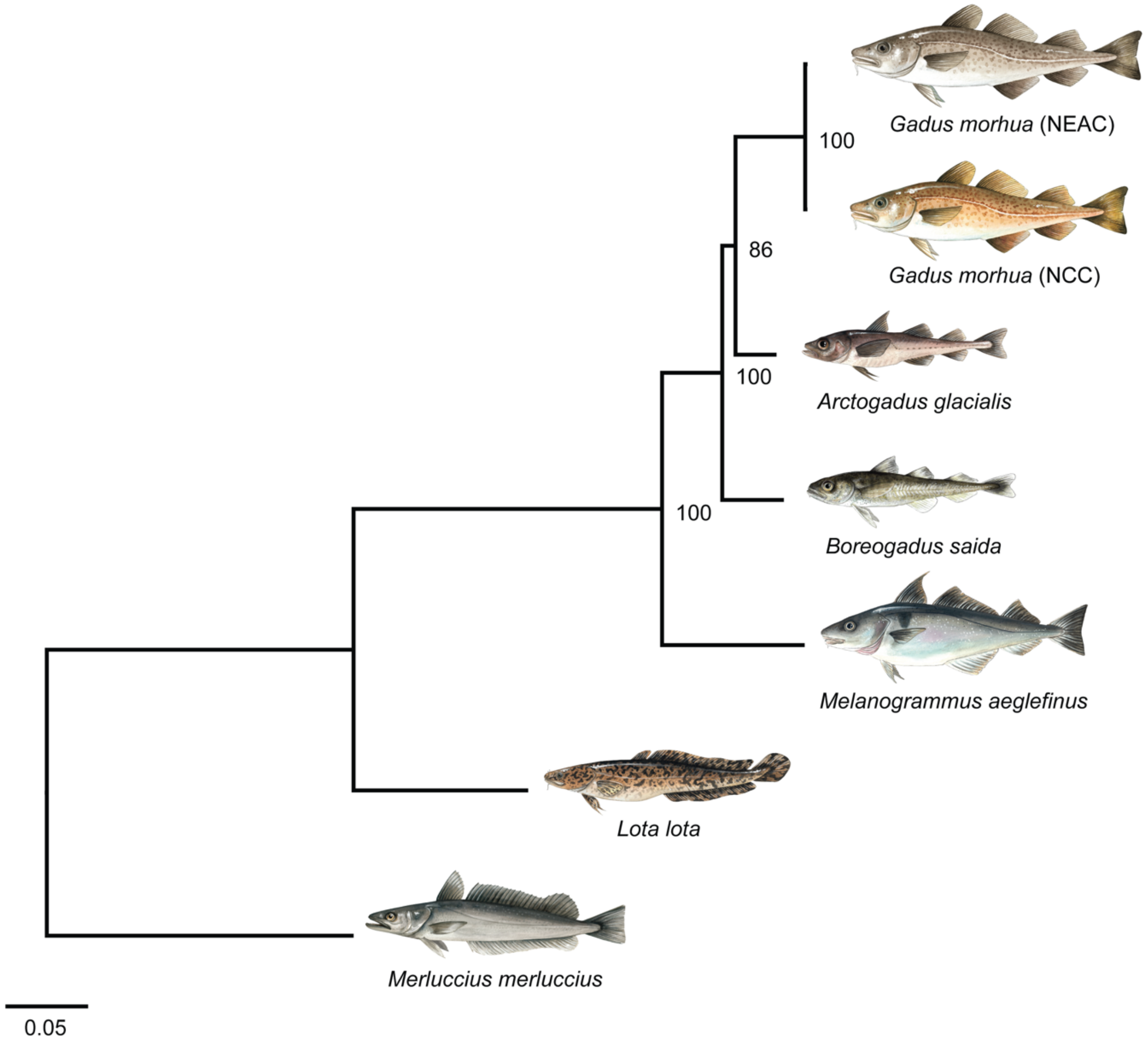

**Figure S11.** Mitochondrial phylogeny of taxa with sequenced genomes in this study, except for European hake which was accessed from NCBI accession: NC_007396.1. ML phylogeny of the focal species using a concatenation approach of all 13 protein-coding genes with maximum likelihood tree inference in IQ-Tree2[55]. Branch length is given in substitutions per site, and bootstrap support is given. All fish illustrations are by Alexandra Viertler.

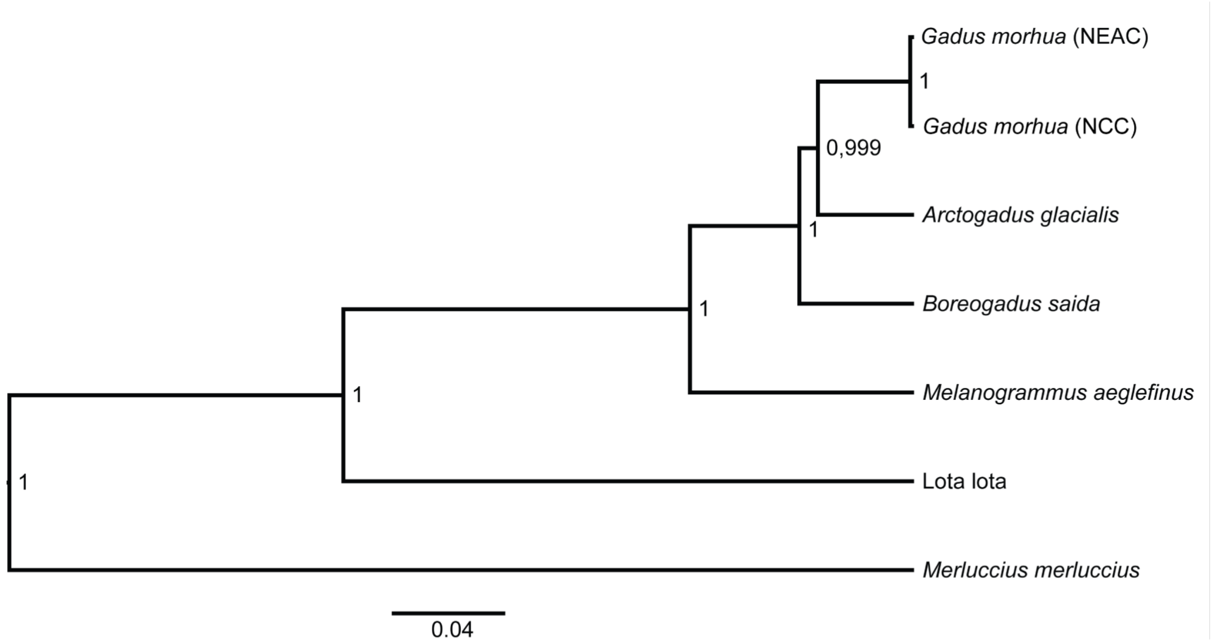

**Figure S12.** Bayesian phylogenetic tree of taxa with sequenced genomes included in this study, except for European hake which was accessed from NCBI accession: NC_007396.1. Bayesian phylogeny of the focal species using a concatenation approach of all 13 protein-coding genes inferred in BEAST2[58]. Branch length is given in substitutions rate, and support given as posterior probabilities.

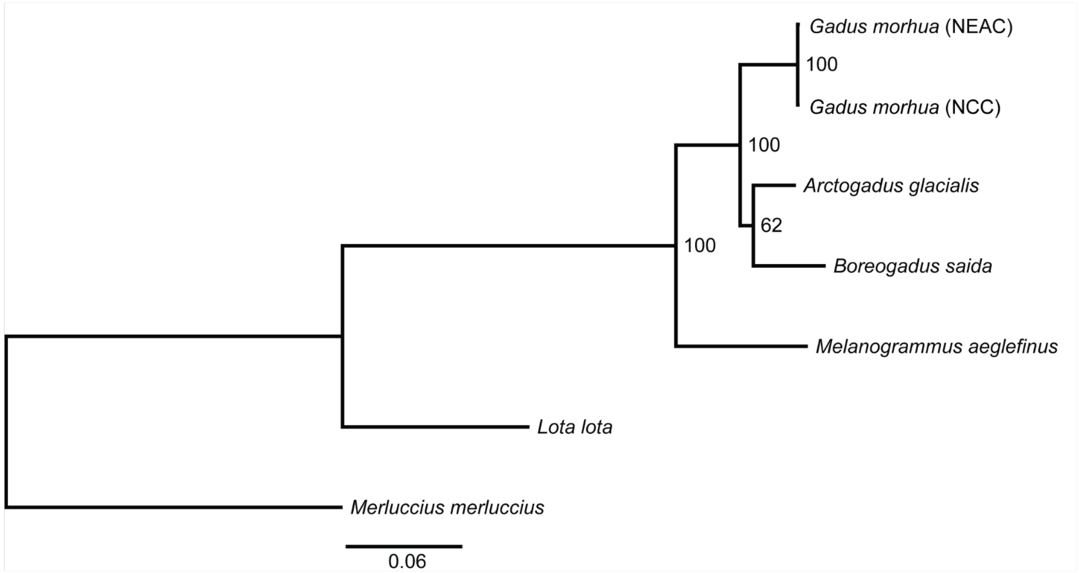

**Figure S13.** Maximum likelihood phylogenetic tree of complete mitogenomes inferred in IQ-Tree2[55]. Bootstrap support shown and branch length given as substitution rate.

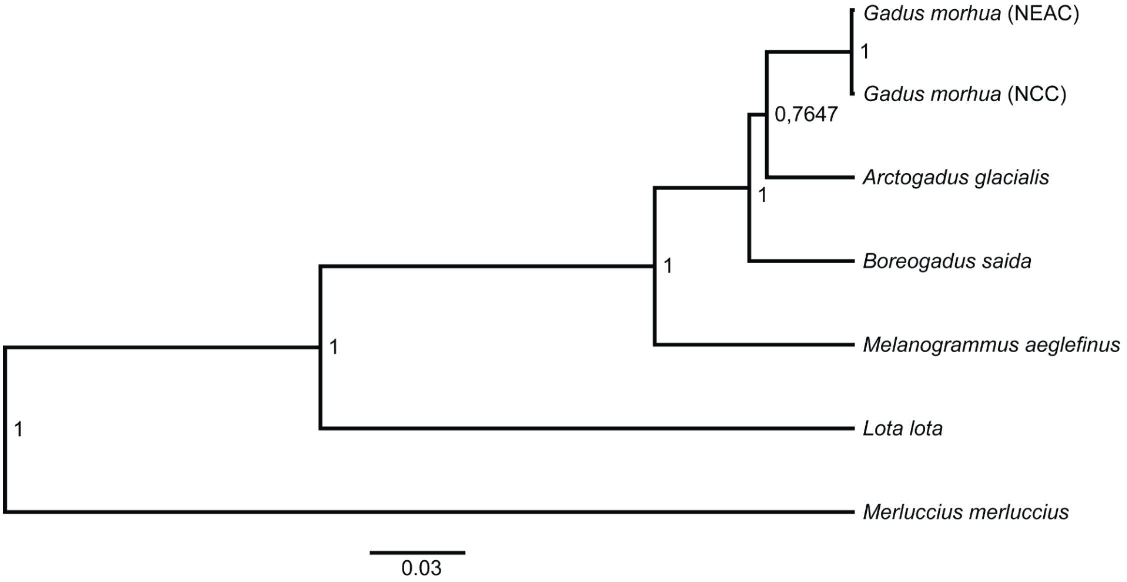

**Figure S14.** Bayesian phylogenetic tree of complete mitogenomes inferred in BEAST2[58], support values given as posterior probabilities and branch length given as substation rate.

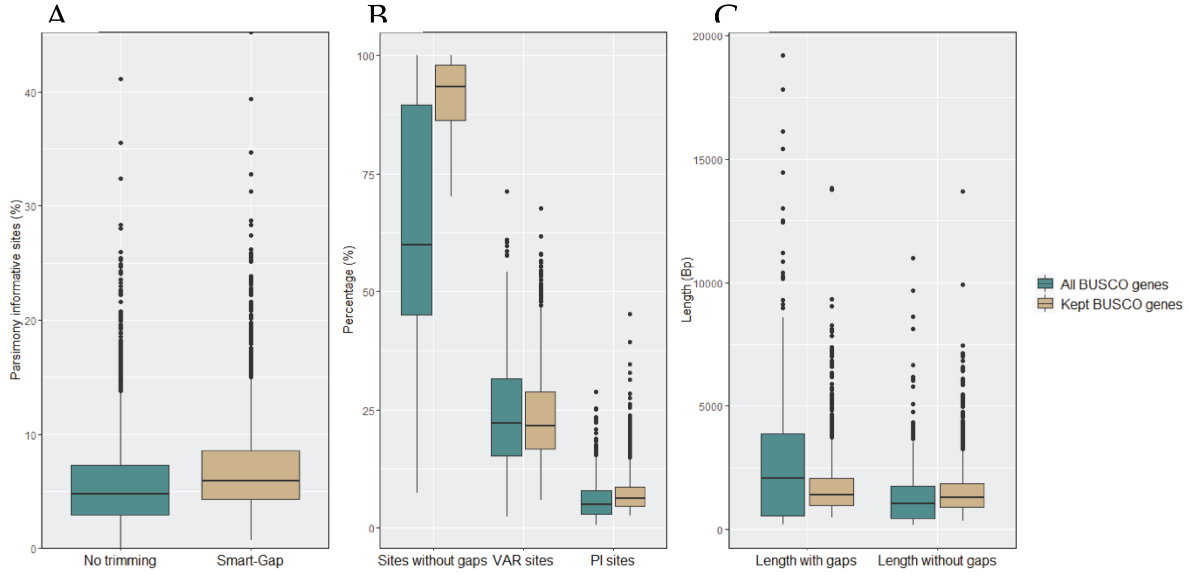

**Figure S15.** Phykit[54] stats represented as boxplot for **A)** percentage of parsimony informative (PI) sites under no trimming versus smart-gap trimming with ClipKIT[61], **B)** sites without gaps, variable (VAR) sites and PI sites after removing BUSCO genes with more than 30% sites containing gaps, less than 5% VAR sites and 2.5% PI sites, **C)** length of BUSCO genes with and without gaps after filtering on minimum length of 500 Bp. Blue box plots show all BUSCO genes and brown box plots show BUSCO genes after filtering.

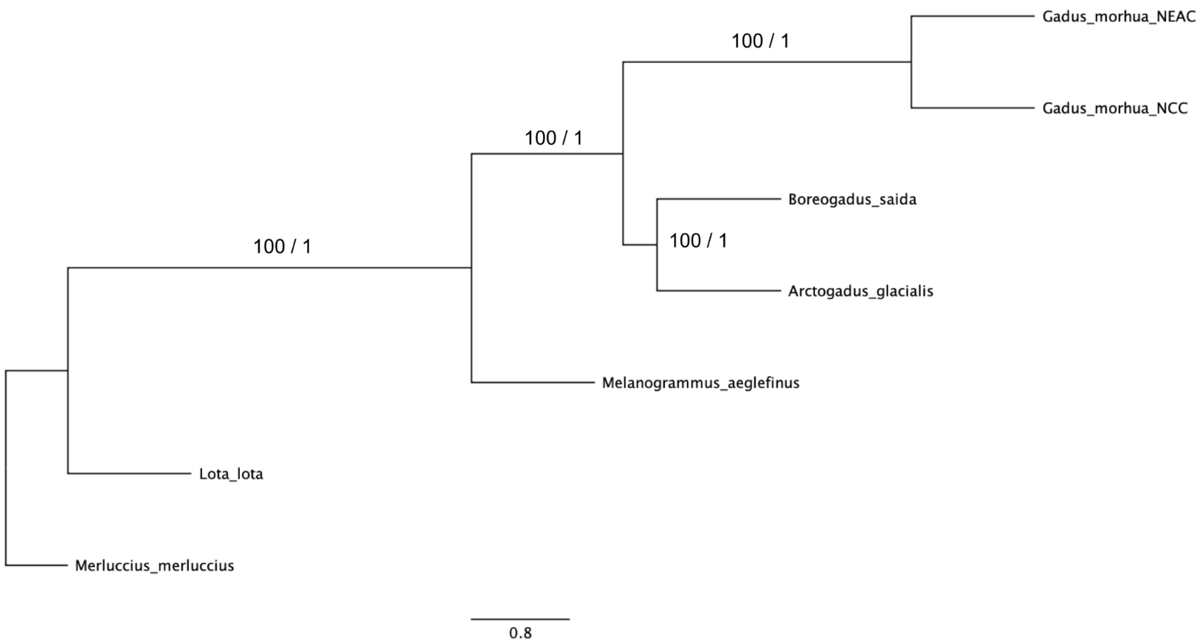

**Figure S16.** Species tree inferred using Astral-III[64] that corresponds to the tree in figure 6. Support values given as bootstrap / posterior probability and branch lengths given as coalescent units.

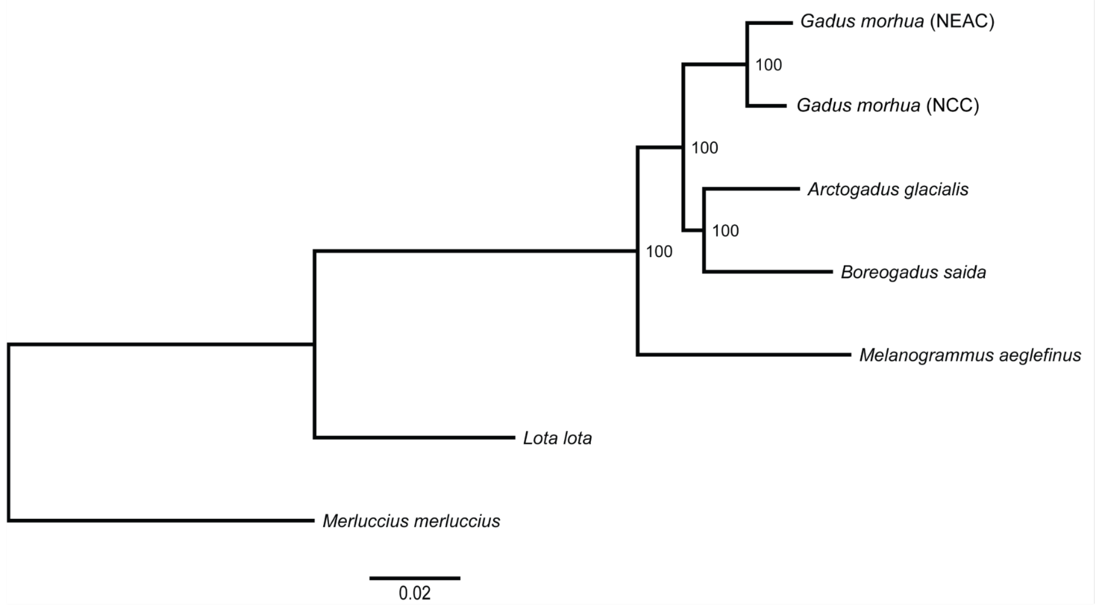

**Figure S17.** Maximum likelihood tree inferred in IQ-Tree2 using 1939 BUSCO genes concatenated into a supermatrix. Bootstrap supports are shown and branch lengths are given in nucleotide substitutions per site.

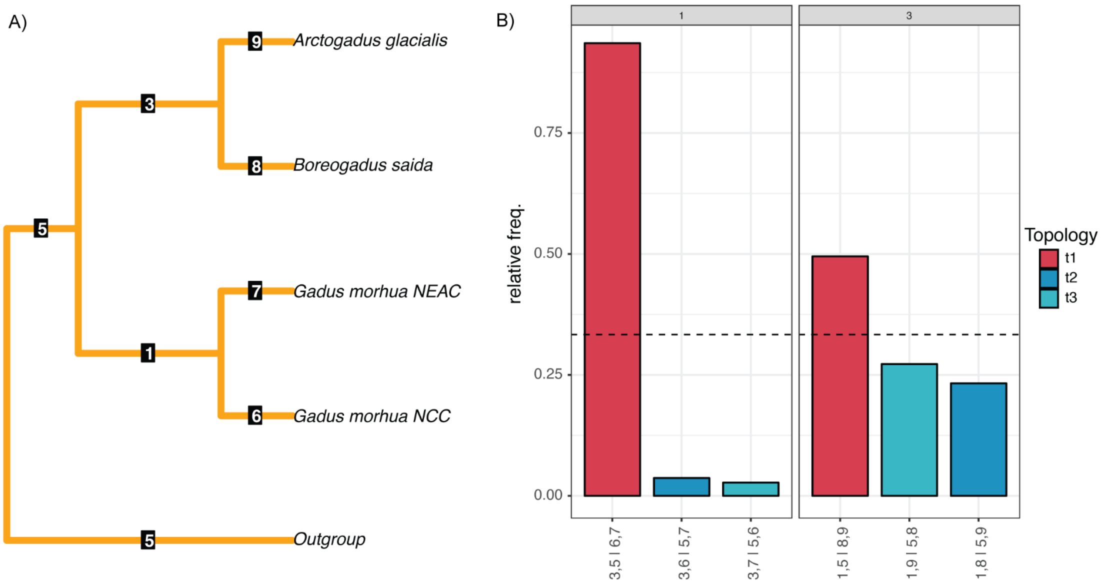

**Figure S18.** The output of quartet frequencies from DiscoVista[65] with *Merluccius merluccius, Lota lota,* and *Melanogrammus aeglefinus* specified as outgroups. **A)** Species tree with internal branch numbers given. **B)** Quartet frequencies are given for each possible topology of the internal branches of interest (1 and 3). Possible topologies are given on the x-axis.

#### Supplementary Tables

**Table S1.** Karyotyping by cytogenetics of Gadiform fishes previously published. 2n=diploid chromosomes reported, and n=inferred haploid chromosomal number. NCBI reference for mitochondrial genes obtained for extended species phylogenetic tree also shown.

| Species | n | 2n | References cytogenetic studies | NCBI accessions for mitogenome: |
|---|---|---|---|---|
| European hake *(Merluccius merluccius)* | 22 | 42 | García-Souto et al. [66] | mitogenome generated in the present study |
| Marbled eel *(Muraenolepis marmoratus)* | 24 | 48 | R. Arai [67] | |
| Polar cod *(Boreogadus saida)* | 18, 19 | 36, 38 | Ghigliotti et al. [68] | mitogenome generated in the present study |
| Arctic cod (*Arctogadus glacialis*) | 14, 15, 16 | 28, 30, 32 | Ghigliotti et al. [68,69] | mitogenome generated in the present study |
| Saffron cod (*Eleginus gracilis)* | 13 | 26 | Ishii K, Yabu H [70] | |
| Navaga *(Eleginus nawaga)* | 13, 13.5 | 26, 27 | Klinkhardt et al. [71] | |
| Pacific cod *(Gadus macrocephalus)* | 22 | 44 | Ishii K, Yabu H [70] | NC_036931.1 |
| Atlantic cod *(Gadus morhua)* | 23 | 46 | Nygren et al. [72], Ghigliotti et al. [73] | mitogenome generated in the present study (NEAC and NCC) |
| Atlantic haddock *(Melanogrammus aeglefinus)* | 22 | 44 | Steve P. Bryson et al. [74] | mitogenome generated in the present study |
| Burbot *(Lota lota)* | 24 | 48 | Lech Kirtiklis et al. [75], Zhou et al. [76] | mitogenome generated in the present study |
| Alaskan pollock *(Gadus chalcogrammus)* | 22 | 44 | Ishii K, Yabu H [70] (Referred to as *Theragra chalcogrammus*) | NC_004449.1 |
| Saithe *(Pollachius virens)* | 20 | 40 | Nygren et al. [72] | NC_015094.1 |
| Blue whiting *(Micromesistius poutassou)* | 22 | 44 | Nygren et al. [72] | NC_015102.1 |
| Pollack *(Pollachius pollachius)* | 19 | 38 | Nygren et al. [72] | NC_015097.1 |
| Poor cod *(Trisopterus minutus)* | 24 | 48 | Nygren et al. [72] | Malmstrøm et al.[5]: utg7180000000029 |
| Forkbeard *(Phycis phycis)* | 24 | 48 | R. Arai [67] | |

**Table S2**. PacBio Sequencing statistics for the six codfishes sequenced.

| Species | Total number of polymerase bases (Gb) | Number of reads | SMRT cells | N50 subread length (kb)* | Average polymerase read length | PacBio sequencing platform | Average subread length (kb) |
|---|---|---|---|---|---|---|---|
| Arctic cod | 43.5 | 6 339 559 | 11 | 9.3 – 10.8 | 5.7 – 8.7 kb | Sequel | 4.5 – 6.9 |
| Polar cod | 47.26 | 6 283 755 | 12 | 9.8 – 11.3 | 6 – 10 kb | Sequel | 4.6 – 7 |
| Atlantic haddock | 98.97 | 5 934 413 | 6 | 17 | 17 kb | Sequel | 12.57 |
| Burbot | 42.42 | 5 414 851 | 10 | 11.3 – 13.3 | 6.9 – 8.6 kb | Sequel | 5.9 – 7.9 |
| European hake | 528 | 9 766 874 | 2 | | 53.7-54.5 kb | Sequel II | |
| Atlantic cod (NCC) Round 1 | 15.80265 | 1 492 072 | 15 | 8 530 – 8 881 | 9 216 - 12 374 bp | RS | |
| Atlantic cod (NCC) Round 2 | 19.24 | 2 395 065 | 8 | | 7475 - 8897 bp | Sequel | |
| Atlantic cod (NCC) Round 3 | 5.46482 | 522 105 | 5 | 10 511 - 10 686 | 10 115 - 10 941 bp | RSII | |

*For Atlantic cod (NCC) N50 insert length (bp) is reported.

**Table S3.** Pairwise comparative syntenic comparison of homologous chromosomes among codfish genome assemblies. Blue shade shows reference species for each comparison.

| Chromosomal number, and the homologous chromosomes between each of the codfish species | | | | | | Decision on most likely event based on reference (blue) vs. queries | Extra comments |
|---|---|---|---|---|---|---|---|
| **Polar cod (Bs)** | **Arctic cod (Ag)** | **Atlantic cod (NEAC) (Ne)** | **Atlantic haddock (Ma)** | **Burbot (Ll)** | **European hake (Mm)** | | |
| 1 | 2*, 4* | 6, 11, 8**, 13** | 5, 8 | 3, 12* | 10, 17 | Fusion in polar cod | |
| 2 | 3*, 5* | 16, 20 | 2, 21 | 8, 14 | 1**, 18**, 3*, 9 | Fusion in polar cod | |
| 3 | 12, 7*,9** | 3, 15 | 1, 6** | 16, 19 | 4, 19 | Fusion in polar cod | Fusion in haddock |
| 4 | 2*, 4* | 08, 22 | 12, 18 | 18, 20 | 6, 18 | Fusion in polar cod | |
| 5 | 6*, 8*, 15** | 13, 17 | 22, 14 | 4, 17 | 1*,2**,5 | Fusion in polar cod | |
| 6 | 9 | 4 | 6 | 1* | 1**, 2*, 15* | | Fission in European hake |
| 7 | 10 | 12 | 4 | 21 | 11 | | |
| 8 | 11 | 7 | 6 | 1**, 15 | 2**, 13 | | Fission in burbot and European hake |
| 9 | 6* | 14 | 7 | 12* | 7 | | |
| 10 | 14 | 10 | 9 | 10 | 8 | | |
| 11 | 1* | 1 | 10 | 17** | 3*, 5** | | Fission in European hake |
| 12 | 13**, 15 | 2 | 13 | 2 | 2*, 6** | | Fission in European hake |
| 13 | 5* | 9 | 11 | 9, 18** | 12 | | Fission in burbot |
| 14 | 13 | 5 | 16 | 6 | 14 | | |
| 15 | 1* | 18 | 17 | 11 | 16 | | |
| 16 | 7* | 21 | 19 | 13 | 21 | | |
| 17 | 8* | 23 | 15 | 7 | 20 | | |
| 18 | 4**, 3* | 19 | 20 | 5 | 1*, 15* | | Fission in European hake |

| | | | | | | |
|---|---|---|---|---|---|---|
| 11, 15, 3** | 1 | 1, 18, 15** | 1**, 4**, 10, 17 | 22, 11 | 3*, 16, 5** | Fusion in Arctic cod |
| 1*, 4* | 2 | 6, 8, 11** | 5, 12 | 20, 3 | 6, 10 | Fusion in Arctic cod |
| 2, 18 | 3 | 16, 19 | 2*, 13**, 20, 21* | 5, 8 | 1*, 15*, 9 | Fusion in Arctic cod |
| 1*, 4* | 4 | 6**, 11, 22 | 8, 18 | 12*, 18 | 17, 18 | Fusion in Arctic cod |
| 2*, 13 | 5 | 9, 20 | 2*, 11, 21* | 14, 9 | 1**, 3*, 12 | Fusion in Arctic cod |
| 5*, 9 | 6 | 14, 17 | 7, 22 | 12*, 4 | 1*, 7 | fusion in Arctic cod |
| 3*, 16 | 7 | 3**, 15, 21 | 1*, 19 | 19, 13, 16 | 21, 4**, 19* | Fusion in Arctic cod |
| 5*, 17 | 8 | 13, 23 | 14, 15 | 7, 17 | 20, 5 | Fusion in Arctic cod |
| 6 | 9 | 4 | 3 | 23, 1* | 1**, 15*, 2* | |
| 7 | 10 | 12 | 4 | 21 | 11 | |
| 8 | 11 | 7 | 6 | 15, 1** | 13, 2** | |
| 3* | 12 | 3, 15** | 1*, 4** | 16, 19** | 4, 19** | Potential translocation of part of Ne15 on to Ne3 resulting in Ag12 |
| 12**, 14 | 13 | 5, 7** | 16, 6** | 6 | 14 | Potential translocation of part of Ne7 on to Ne5 resulting in Ag13 |
| 10 | 14 | 10 | 9 | 10 | 8 | |
| 5**, 12 | 15 | 2 | 13 | 2 | 2* | |
| 11 | 1* | 1 | 10 | 17**, 22 | 3*, 5** | |
| 12 | 15 | 2 | 13 | 2 | 2*, 6** | |
| 3* | 12, 7** | 3 | 1* | 16 | 4 | |
| 6 | 9 | 4 | 3 | 1*, 23 | 1**, 2*,15* | |
| 14 | 13 | 5 | 16 | 6 | 14 | |
| 1* | 2*, 4** | 6 | 5, 8** | 3, 12** | 10, 17** | |
| 8 | 11 | 7 | 6 | 1**, 15 | 2**, 13 | |
| 4* | 2* | 8 | 12 | 20 | 6 | |
| 13 | 5* | 9 | 11 | 9 | 12 | |
| 10 | 14 | 10 | 9 | 10 | 8 | |
| 1* | 4*, 2** | 11 | 5*, 8 | 3**, 12* | 10**, 17 | |
| 7 | 10 | 12 | 4 | 21 | 11 | |
| 5* | 8* | 13 | 14 | 17 | 5 | |
| 9 | 6* | 14 | 7 | 12* | 7 | |
| 3* | 7*, 12**, 1** | 15 | 1* | 19 | 19 | |
| 2* | 3* | 16 | 2*, 21* | 8 | 9 | |
| 5* | 6* | 17 | 22 | 4 | 1* | |
| 15 | 1* | 18 | 17 | 11 | 16 | |

| | | | | | |
|---|---|---|---|---|---|
| 18 | 3* | 19 | 20 | 5 | 1*, 15* |
| 2* | 5* | 20 | 2*, 21* | 14 | 1**, 3* |
| 16 | 7* | 21 | 19 | 13 | 21 |
| 4* | 4* | 22 | 18 | 18 | 18 |
| 17 | 8* | 23 | 15 | 7 | 20 |

*Makes up half of the chromosome in the query.
**Small homologous chromosomal regions likely make up inter-chromosomal translocations or multiple hits (i.e., duplicated genes).

**Table S4.** BUSCO gene completeness statistics across final genome assemblies for six species, and the Atlantic cod (NEAC) reference genome gadMor3[4].

| Species | Complete / total searched | CS | CD | Fragmented | Missing |
|---|---|---|---|---|---|
| Arctic cod | 3337 / 3640 | 3294 / 90.5% | 43 / 1.2% | 53 / 1.5% | 250 / 6.8% |
| Polar cod | 3282 / 3640 | 3234 / 88.8% | 48 / 1.3% | 54 / 1.5% | 304 / 8.4% |
| Atlantic cod (NCC) | 3342 / 3640 | 3294 / 90.5% | 48 / 1.3% | 33 / 0.9% | 265 / 7.3% |
| Atlantic cod (NEAC) (gadMor3.0) | 3404 / 3640 | 3342 / 91.8% | 62 / 1.7% | 27 / 0.7% | 209 / 5.8% |
| Atlantic haddock (ENA: ERR1473879) | 3291 / 3640 | 3234 / 88.8% | 57 / 1.6% | 54 / 1.5% | 295 / 8.1% |
| Atlantic haddock (present study assembly) | 3427 / 3640 | 3427 / 94.1% | 33 / 0.9% | 63 / 1.7 % | 150 / 4.2% |
| Burbot | 3455 / 3640 | 3418 / 93.9% | 37 / 1.0% | 22 / 0.6% | 163 / 4.5% |
| European hake | 3324 / 3640 | 3216 / 88.4% | 108 / 3.0% | 29 / 0.8 % | 287 / 7.8% |

CS: Complete and single-copy, CD: Complete and duplicated.
Total BUSCO groups searched: 3640.

**Table S5.** Specimens sequenced for genome assemblies in the present study.
* Denotes tissue stored on 96% EtOH, ** denotes sex i.e., M(Male) or F(Female).

| Specimen | Sample date | Origin | Weight | Length | Sex** / Age (years) | Tissue (PacBio / Illumina) | Tissue (10X) | Tissue (Hi-C) |
|---|---|---|---|---|---|---|---|---|
| Arctic cod *(Arctogadus glacialis)* | 13/08/ 2013 | Tyroler Fjord, Greenland | 79.0 g | 23.4 cm | F / 6 | Gill filaments* | Gill filaments* | Gill filaments* |
| Polar cod *(Boreogadus saida)* | 13/08/ 2013 | Tyroler Fjord, Greenland | 100.7 g | 23.6 cm | M / 6 | Spleen/ liver/ gill* | Gill filaments* | Gill filaments* |
| Norwegian coastal cod (NCC) *(Gadus morhua)* | 29/06/ 2011 | Lofoten, Norway | 1600.0 g | 62.0 cm | F / 7 | Spleen* | Spleen** | Spleen** |
| Atlantic haddock *(Melanogrammus aeglefinus)* | 23/10/ 2019 | Polaria Aquarium, Tromsø, Norway | 780.0 g | 44.0 cm | M | Blood* | Blood* | Blood* |
| Burbot *(Lota lota)* | 06/09/ 2017 | Stuorajávri, Kautokeino, Norway | 743.1 g | 49.0 cm | F | Spleen/ gill/ liver* | | Gill* |
| European hake *(Merluccius merluccius)* | 10/05/ 2021 | Inner Oslofjord, Norway | | | M | Blood* | | Blood* |

**Table S6.** Overview and metadata parameters of population samples of Arctic cod and polar cod used in the present study for genetic differentiation estimates. For more information on sequencing of samples see Hoff et al.[77] for polar cod and Maurstad et al.[78] for Arctic cod.

| Sample ID | Length (mm) | Weight (g) | Sex | Maturity | Trawling location |
|---|---|---|---|---|---|
| 01-Ag13010 | 206 | 72 | M | 1 | Greenland, Tyroler |
| 02-Ag13014 | 189 | 49.9 | F | 1 | Greenland, Tyroler |
| 03-Ag13019 | 266 | 133 | F | 2 | Greenland, Tyroler |
| 04-Ag13020 | 203 | 56.6 | F | 1 | Greenland, Tyroler |
| 05-Ag13021 | 231 | 82.9 | F | 1 | Greenland, Tyroler |
| 06-Ag13024 | 210 | 67.4 | F | 1 | Greenland, Tyroler |
| 07-Ag13026 | 182 | 45 | F | 0 | Greenland, Tyroler |
| 08-Ag13027 | 204 | 70.2 | M | 1 | Greenland, Tyroler |
| 09-Agl17001 | 203 | 59.55 | F | 1 | Greenland, Besselfjord |
| 10-Agl17002 | 181 | 39.6 | M | 1 | Greenland, Besselfjord |
| 11-Agl17003 | 155 | 21.5 | * | * | Greenland, Besselfjord |
| 12-Ag-ds-0A08045-2 | * | * | * | * | Davis Strait |
| 13-Ag1-hyb-aen-2018 | 131 | 16 | F | 1 | Barents Sea |
| SAMEA4028798 | * | * | * | * | Davis Strait |
| 04-polcod59 | 15.5 | 24 | F | 2 | Barents Sea |
| 171-polcod1 | 15 | 22.75 | M | 2 | Barents Sea |
| 172-polcod2 | 14.1 | 19.35 | M | 2 | Barents Sea |
| 174-polcod4 | 13.6 | 18.75 | F | 2 | Barents Sea |
| 178-polcod8 | 15.7 | 24.45 | M | 4 | Barents Sea |
| 185-polcod20 | 17 | 34 | M | 2 | Barents Sea |
| 187-polcod37 | 14.3 | 22 | M | 2 | Barents Sea |
| 188-polcod45 | 17.5 | 35 | F | 3 | Barents Sea |
| 189-polcod49 | 13.3 | 18 | F | 2 | Barents Sea |
| 190-polcod50 | 16 | 35 | F | 2 | Barents Sea |
| 192-polcod52 | 19,5 | 55 | F | 3 | Barents Sea |
| 228-polcod62 | 18.4 | 49 | M | 4 | Barents Sea |
| 235-polcod74 | 18.8 | 48 | M | 3 | Barents Sea |
| 287-polcod31 | 18 | 39 | F | 3 | Barents Sea |

*No information available. +Accessed from ENA: ERR1473882, ERR1473883.
